# The quantitative genetics of the prevalence of infectious diseases: hidden genetic variation due to Indirect Genetic Effects dominates heritable variation and response to selection

**DOI:** 10.1101/2021.04.07.438789

**Authors:** Piter Bijma, Andries D. Hulst, Mart C. M. de Jong

## Abstract

Infectious diseases have profound effects on life, both in nature and agriculture. Despite the availability of well-established epidemiological theory, however, a quantitative genetic theory of the host population for the endemic prevalence of infectious diseases is almost entirely lacking. While several studies have demonstrated the relevance of the transmission dynamics of infectious diseases for heritable variation and response to selection of the host population, our current theoretical framework of quantitative genetics excludes these dynamics. As a consequence, we do not know which genetic effects of the host population determine the prevalence of an infection, and have no concepts of breeding value and heritable variation for endemic prevalence.

Here we integrate quantitative genetics and epidemiology, and propose a quantitative genetic theory for *R*_*0*_ and for the endemic prevalence of an infectious disease. We first identify the genetic factors that determine the prevalence of an infection, using an approach founded in epidemiological theory. Subsequently we investigate the population level consequences of individual genetic variation, both for *R*_0_ and for the endemic prevalence. Next, we present expressions for the breeding value and heritable variation, for both prevalence and individual binary disease status, and show that these parameters depend strongly on the level of the prevalence. Results show that heritable variation for endemic prevalence is substantially greater than currently believed, and increases when prevalence approaches zero, while heritability of individual disease status goes to zero. As a consequence, response of prevalence to selection accelerates considerably when prevalence goes down, in contrast to predictions from classical theory. Finally, we show that most of the heritable variation for the endemic prevalence of an infection is hidden due to indirect genetic effects, suggesting a key role for kin-group selection both in the evolutionary history of current populations and for genetic improvement strategies in animals and plants.

## Introduction

Pathogens have profound effects on life on earth, both in nature and agriculture, and also directly on the human population (Schrag and Wiener, 1995; Russel, 2013). In nature, infectious pathogens are a major force shaping evolution of populations by natural selection, both in animals and plants (reviewed in Karlsson *et al*. 2014 and Ebert and Fields 2020). In livestock, the annual cost of fighting and controlling epidemic and endemic infectious diseases is substantial, and much greater than the annual value of genetic improvement (Rushton, 2009; Knap and Doeschl-Wilson, 2020). Moreover, while antimicrobials have revolutionized medicine, the rapid appearance of resistant strains has resulted in a global health problem, both in the human population and in livestock (EFSA 2012; Thanner *et al*. 2016). Thus there is an urgent need for additional methods and tools to combat infectious diseases. For livestock and plant production, artificial genetic selection of (host) populations for infectious disease traits may provide such a tool. To quantify and optimize the potential benefits of such selection, however, we need to understand the quantitative genetics of infectious disease traits.

Despite the availability of well-established epidemiological theory (*e*.*g*., Diekmann *et al*. 2012), a quantitative genetic theory of the host population for the prevalence of infectious diseases is almost entirely lacking. Current approaches for genetic selection against infectious diseases in livestock and crops are entirely based on the individual host response, ignoring transmission of the infection in the population. While several studies have demonstrated the relevance of the transmission dynamics of infectious diseases for heritable variation and response to selection in the host population (Lipschutz-Powell *et al*., 2012; Anche *et al*., 2014; Tsairidou *et al*., 2019; Hulst *et al*., 2021), mostly using stochastic simulations, the current theoretical framework of quantitative genetics does not include these dynamics. Moreover, we lack general expressions for the genetic variance in key epidemiological parameters, in particular the basic reproduction number *R*_0_, even though such parameters may have a genetic basis.

Infections for which recovery does not confer any long-lasting immunity typically show endemic behaviour, where the infection remains present in the population. For such infections, the endemic prevalence is defined as the expected fraction of the population that is infected. Because we lack a theoretical quantitative genetic framework for infectious diseases, we do not know which genetic effects of the host population determine the prevalence of an infectious disease, and have no concepts of breeding value and heritable variation for endemic prevalence. Hence, we do not understand response to genetic selection in the endemic prevalence of infectious diseases at present. The main parameter determining the prevalence of endemic infections is the basic reproduction number *R*_0_, defined as the average number of individuals that gets infected by a typical infected individual in an otherwise non-infected population. In this manuscript we will propose a quantitative genetic framework for heritable variation and response to selection for *R*_0_ and for the endemic prevalence of infectious diseases.

Individual phenotypes for infectious diseases are often recorded as the binary infection status of an individual, zero indicating non-infected and one indicating infected. The prevalence of an infection is then defined as the fraction of individuals that is infected, which is the fraction of individuals that has infection status *y* = 1. Because the average value of individual binary infection status is equal to the fraction of individuals infected, response to genetic selection in binary infection status is identical to response in prevalence, and *vice versa*. Binary infection status (0/1) typically shows low heritability, which suggests that response to selection is limited, also for prevalence (Bishop and Woolliams 2010; Bishop *et al*., 2012; Martin *et al*., 2018).

Geneticists have long realized that the categorical distribution of binary traits does not agree well with quantitative genetic models for polygenic traits, such as the infinitesimal model (Fisher 1918). For this reason, models have been developed that link an underlying normally-distributed trait to the observed binary phenotype, such as the threshold model (Dempster & Lerner, 1950; Gianola, 1982) and the equivalent generalized linear mixed model with a probit link function (*e*.*g*., de Villemereuil *et al*. 2016). In such models, the underlying scale is interpreted as causal, and genetic parameters are assumed to represent “biological constants” on this scale. The genetic parameters on the observed scale, in contrast, depend on the mean of the trait, and thus change with the mean even when the change in allele frequencies at causal loci is infinitesimally small. In a landmark paper, Robertson (1950) showed that the observed-scale heritability of binary traits reaches a maximum at a prevalence of 0.5, and approaches zero when the prevalence is close to 0 or 1. Hence, observed-scale heritability vanishes when artificial selection moves prevalence close to zero, hampering further genetic change.

Infectious disease status, however, differs fundamentally from binary phenotypes for non-communicable traits, such as, say, heart failure. Because pathogens can be transmitted between host individuals, either directly or via the environment, the infection status of an individual depends on the status of other individuals in the population. This suggests that Indirect Genetic Effects (IGE) may play a role, which would fundamentally alter heritable variation and response to selection (Griffing 1967; Moore *et al*., 1997; Wolf *et al*., 1998; Bijma and Wade 2008; Bijma 2011). Results of simulation studies indeed suggest that selection response in the prevalence of infectious diseases may differ qualitatively from response in non-communicable traits (Nieuwhof *et al*., 2009; Doeschl-Wilson *et al*., 2011; Anche *et al*., 2014; Hulst *et al*. 2021), and this has also been observed in an actual population (Heringstad *et al*. 2007). Results of Hulst *et al*. (2021), for example, show that genetic selection may result in the eradication of an infection via the mechanism of herd immunity, just like with vaccination (FINE 1993). This result contradicts predictions based on the observed-scale heritability for non-communicable binary traits, where heritability vanishes when prevalence approaches zero (Robertson, 1950).

While quantitative geneticists and breeders typically focus on individual disease status and (implicitly) interpret prevalence as an average of individual trait values, epidemiologists interpret the endemic prevalence of an infectious disease as the result of a population level process of transmission of the infection (Kermack and McKendrick, 1927; Keeling and Rohani, 2011; Diekmann *et al*. 2012). In the latter perspective, both *R*_0_ and the prevalence are emergent properties of a population, similar to the size of a termite colony or the number of prey caught by a hunting pack, rather than an average of individual trait values. Because such emergent traits do not belong to single individuals, we cannot apply the common partitioning of individual phenotypic values into individual additive genetic values (breeding values) and non-heritable residuals (“environment”). Nevertheless, the genetic effects that determine the response to selection in an emergent trait and the heritable variation for an emergent trait can be defined based on the so-called total heritable variation (Bijma 2011). The total heritable variation in a trait is based on the individual genetic effects on the level of the emergent trait, rather than on a decomposition of individual trait values into genetic and residual effects. This suggests we can develop a quantitative genetic theory for the endemic prevalence of infectious diseases by combining epidemiological theory with the theory of total heritable variation.

Here we propose a quantitative genetic theory for the basic reproduction number *R*_0_ and for the endemic prevalence of infectious diseases. We first identify the genetic factors that determine the prevalence of an infectious disease. Similar to the threshold model, we will assume an underlying additive infinitesimal model for those genetic factors. However, the link between the underlying additive scale and the observed endemic prevalence will be founded in epidemiological theory, with a key role for *R*_0_. Subsequently we investigate the population level consequences of genetic variation in individual disease traits for *R*_0_ and for the endemic prevalence. Next, we move to the individual level, and derive expressions for the breeding value and heritable variation, for *R*_0_, endemic prevalence and individual binary infection status, and show how these parameters depend on the level of the endemic prevalence. Results will show that heritable variation for endemic prevalence increases when prevalence approaches zero, while heritability of individual infection status goes to zero. Then we investigate response to selection, and show that response of prevalence to selection accelerates considerably when prevalence goes down. Finally, we partition the breeding value for prevalence into direct and indirect genetic effects, and show that most of the heritable variation in the endemic prevalence of the infection is indirect, and thus hidden to classical genetic analysis and selection. We focus solely on the development of quantitative genetic theory, and do not consider the statistical estimation of the genetic effects underlying prevalence. Such methods have been developed elsewhere (Anacleto *et al*. 2015; Biemans *et al*. 2017; Pooley *et al*. 2020).

## Theory and Results

### 1. The genetic factors that determine R_0_ and the endemic prevalence

We consider an endemic infectious disease, where individuals can either be susceptible (*i*.*e*., in the non-infected state), denoted by S, or in the infected state, denoted by I. We use corresponding symbols in *italics* to denote the number of individuals with that status. Thus, with a total of *N* individuals in the population in which the endemic takes place, *S* denotes the number of susceptible individuals, *I* the number of infected individuals, and *S* + *I* = *N* (see Table 1 for a notation key). We will assume that infected individuals are also infectious, and can thus infect others. When individuals recover they become susceptible again. This model is known as the SIS compartmental model (Hethcote, 1989), and was first discussed by Weiss and Dishon (1971; In the Discussion we will consider the validity of our results for other compartmental models).

**Table 1.**
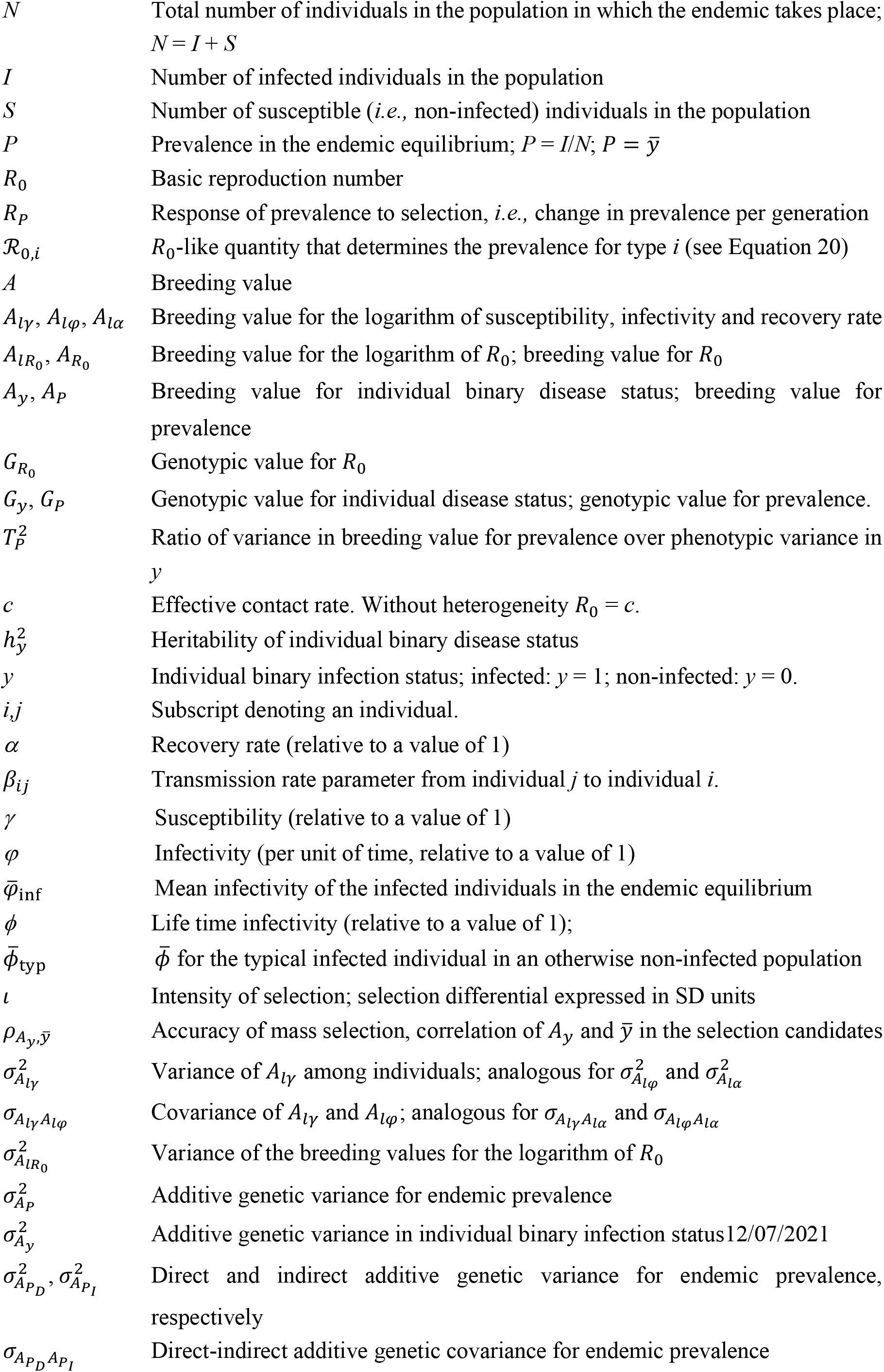
Notation key.

The prevalence (*P*) of an endemic infection is defined as the fraction of the population infected (Diekmann *et al*. 2012),

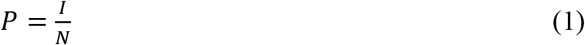

When individual infection status is coded in a binary fashion, using *y* = 0 for non-infected individuals and *y* = 1 for infected individuals, the prevalence is also equal to the average individual infection status in the population,

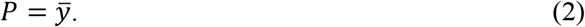

The prevalence of an infectious disease is determined by *R*_0_. The *R*_0_ is defined as the average number of individuals that get infected by a typical (*i*.*e*., average) infected individual in an otherwise non-infected population, and is a property of the population (Kermack and McKendrick, 1927; Anderson and May, 1979; Diekmann *et al*. 1990). When *R*_*0*_ > 1, an average infected individual on average infects more than one new individual in an infection free population, and the infection can persist in the population.

The prevalence of an endemic infection reaches an equilibrium value, known as the endemic prevalence, when a single typical infected individual on average infects one other individual (*R* = 1; the endemic steady state). In a population where all individuals are the same, *i*.*e*., in the absence of genetic heterogeneity among host individuals, this occurs when the product of *R*_0_ and the fraction of contact individuals that is susceptible is equal to one; *R*_0_ (1 − *P*) = 1. For example, when *R*_*0*_ = 3, an infected individual could in principle infect three other individuals. However, when only one third of its contact individuals is susceptible (*i*.*e*., not infected), meaning 1−*P* = 1/3, then the effective reproduction number equals 3 × 1/3 = 1. Hence, when 1−*P* = 1/3, an infected individual is on average replaced by a single newly infected individual, so that an equilibrium occurs at *P* = 1 − 1/3 = 2/3. In the absence of heterogeneity, therefore, the endemic prevalence is given by (Weiss and Dishon 1971).

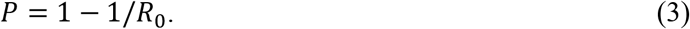

Throughout, we will use the symbol *P* to denote the endemic prevalence. The actual prevalence tends to fluctuate around the equilibrium value because of random perturbations and transient effects, for example when new animals replace some of the resident animals. Equation 3 is an approximation when there is variation among individuals, which is commonly referred to as “heterogeneity” in the epidemiological literature, and which will be addressed in section 4 below.

Figure 1 illustrates the relationship between the endemic prevalence and *R*_0_. When *R*_0_ is smaller than one the endemic prevalence is zero (the infection is not present in the long run), and Equation 3 does not apply. For large *R*_0_ the endemic prevalence asymptotes to 1. This threshold phenomenon, *i*.*e*., *P* = 0 when *R*_0_ < 1 and *P* > 0 when *R*_0_ ≥ 1, is exact also with heterogeneity (Diekmann *et al*., 1990). Note that the curve is steeper the closer *R*_0_ is to 1. This pattern will have considerable consequences for the relationship between the heritable variation in the endemic prevalence and the level of the endemic prevalence, as will be shown in section 6 of this manuscript.

**Figure 1.**
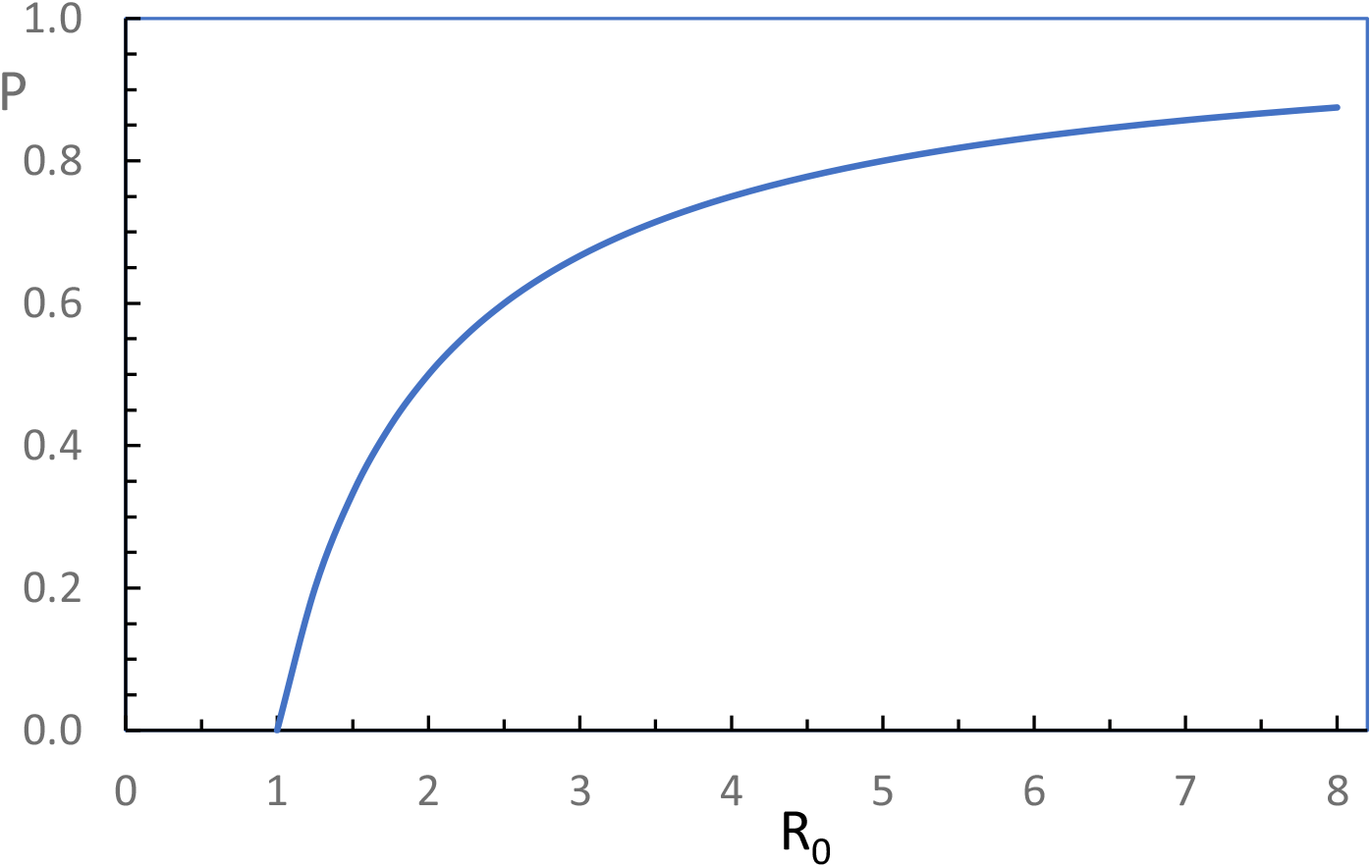
The relationship between the endemic prevalence (*P*) and the basic reproduction number (*R*_0_) for a homogeneous population. From Equation 3.

Because the endemic prevalence is determined by *R*_0_ (Equation 3), the response of prevalence to selection, *i*.*e*., the genetic change in the endemic prevalence from one (host) generation to the next, follows from the genetic change in *R*_0_. Thus, to measure the value of an individual with respect to response to selection, we should base this measure on the genetic impact of the individual on *R*_*0*_. In other words, the definition of an individual breeding value for endemic prevalence should be based on *R*_0_. The next step, therefore, is to find the individual genetic factors underlying *R*_0_.

In the absence of variation among individuals (heterogeneity), *R*_0_ is the product of the transmission rate parameter (*β*) and the mean duration of the infectious period (1/*α*; Kermack and McKendrick, 1927; Diekmann *et al*., 1990),

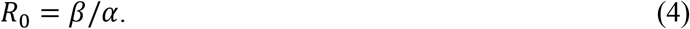

where *α* is the recovery rate parameter. The *β* is the average number of individuals infected per unit of time by a single infected individual when all its contact individuals are susceptible, and α is the probability per unit of time for an infected individual to recover. With heterogeneity, Equation 4 is an approximation (Diekmann *et al*., 1990); the effect of heterogeneity on *R*_0_ will be addressed in section 3 of this manuscript.

With heterogeneity, the transmission rate parameter may vary between pairings (contacts) of individuals. The transmission rate parameter between infectious individual *j* and susceptible individual *i* may be modelled as the product of an overall effective contact rate (*c*), the susceptibility (*γ*) of recipient individual *i* and the infectivity (*φ*) of donor individual *j* (*e*.*g*., De Jong *et al*., 1996; Lipschutz-Powell *et al*., 2014; Anacleto *et al*. 2015; Biemans *et al*. 2017),

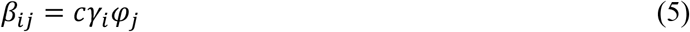

Hence, *β*_*ij*_ refers to transmission from individual *j* to *i*, and may differ from *β*_*ji*_. Note that *i* and *j* are distinct individuals in Equation 5, so that *γ*_*i*_ and *φ*_*j*_ are independent when *i* and *j* are genetically unrelated.

Equations 3 through 5 show that the factors underlying the endemic prevalence of an infection are the contact rate *c*, the susceptibility, *γ*, the infectivity, *φ*, and the recovery rate *α*. We define *c* as a fixed parameter for the population (or, for example, for a sex, herd or age class combination), whereas *γ, φ* and *α* may show random variation among individuals. Note, while the actual contact rate may vary among individuals, it is convenient to include such variation in the individual susceptibility and infectivity traits. Thus we also assume that all the individuals are mixing randomly within the (local) population, such as a herd. In principle, *β*_*ij*_ might also depend on the specific combination of *i* and *j*, so that we cannot separate *β*_*ij*_ into a product of components due to *i* an *j*. However, in a quantitative genetic perspective, such an interaction represents between-individual epistasis, which does not contribute to the heritable variation, and which we will therefore ignore (In epidemiological terminology, we assume separable mixing; Diekmann *et al*. 1990). Hence, conceptually we define *c* as the average effective contact rate for the population, while variation in contact rate among individuals is included in *γ* and *φ*. Moreover, to define the scale of Equations 4 and 5, it is convenient to include the scale in *c*, and to express *γ, φ* and *α* relative to a value of 1. Hence, with this parameterisation, the *c* is on the scale of *R*_0_, and *R*_0_ and *c* are identical in the absence of heterogeneity. With heterogeneity, however, *R*_0_ may deviate from *c* (See section 3 below).

### 2. Genetic models for susceptibility, infectivity, recovery and *R*_*0*_

Genetic variation is potentially present in susceptibility, infectivity and the recovery rate. In this section we propose a genetic model for these traits, which subsequently leads to a genetic model for *R*_0_.

We assume that susceptibility, infectivity and the recovery rate are affected by a large number of loci, each of small effect, so that genetic effects approximately follow a Normal distribution. However, as *γ, φ*, and α represent rates, *i*.*e*., probabilities per unit of time, their values are strictly positive. Moreover, in the expressions for the epidemiological parameters of the previous section (Equations 4 and 5), all these parameters appear in products. For these reasons, following Anacleto *et al*. (2015), we define normally distributed additive genetic effects for the logarithm of these rates, so that effects are multiplicative on the actual scale, and the rates themselves follow a log-normal distribution,

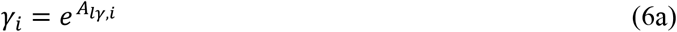

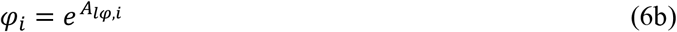

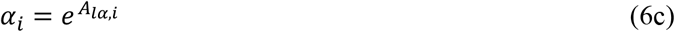

where *A*_*l*..,*i*_ denotes the Normally distributed additive genetic value (breeding value) for the logarithm of the corresponding rate for individual *i*, and has a mean of zero,

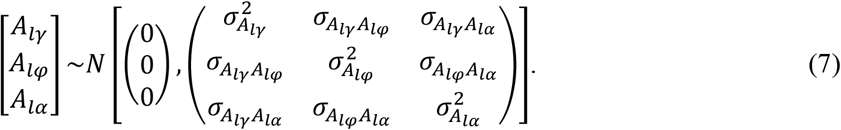

Throughout, we use subscript *l* to denote the natural logarithm. Thus, the breeding values for log (*γ*), log (*φ*) and log (α) follow a multivariate normal distribution, as common in quantitative genetics. Moreover, for the average individual the *A*_*l*.._ = 0, so that its rates are equal to one (*γ* = *φ* = α = 1). Hence, those rates should be interpreted relative to a value of 1. An individual with *γ* = 2, for example, is twice as susceptible as the average individual.

The breeding values on the log-scale can approximately be interpreted as a relative change of the corresponding rate. For example, since *e*^0.1^ *≈* 1.1, an *A*_*lγ*_ of 0.1 corresponds approximately to a 10% greater than average susceptibility (*γ ≈* 1.1). Similarly, an *A*_*lγ*_ of −0.1 corresponds approximately to a 10% smaller than average susceptibility (*γ ≈* 0.9). Realistic values for the genetic variances on the log-scale are probably smaller than ∼0.5^2^ (Hulst *et al*., 2021). For example, with 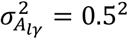, the 10% least susceptible individuals have 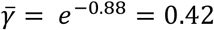, while the 10% most susceptible individuals have 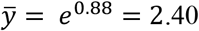. Thus the average susceptibilities of these top and bottom 10% of individuals differ by a factor of 5.7, which is substantial. Therefore, we will consider additive genetic variances on the log-scale no greater than 0.5^2^. With a prevalence of 0.3, this value corresponds to an observed-scale heritability of individual binary infection status of about 0.05 (Hulst *et al*., 2021).

#### Genotypic value and breeding value for *R*_*0*_

Based on Equations 4 and 5, we may define an individual genotypic value for *R*_0_ (see also Anche *et al*. 2014 and Biemans *et al*. 2019),

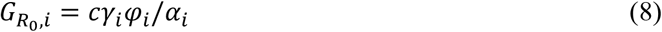

In contrast to the pair-wise transmission rate parameter *β*_*ij*_ in Equation 5, an individual’s genotypic value for *R*_0_ is entirely a function of its own rates, as can be seen from the index *i* on all elements of Equation 8. This is because 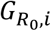 refers to the genetic effects that originate from the individual, rather than to those that affect its trait value. As a consequence these rates may be correlated, as defined in Equation 7 above. Hence, 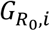 represents a total genotypic value, including both direct and indirect genetic effects (Bijma *et al*., 2007; Bijma 2011). We focus on the total genotypic value, because our ultimate interest is in response to selection (Section 7). In section 3 of this manuscript, we will show that *R*_0_ is indeed the simple population average of 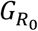.

From Equations 6 and 8,

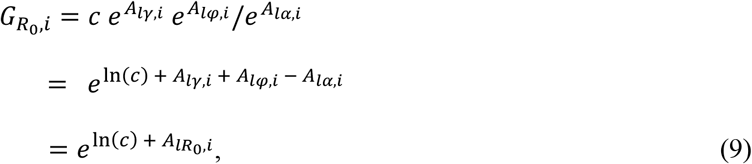

where 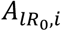 is a Normally distributed additive genetic effect (breeding value) for the logarithm of *R*_0_,

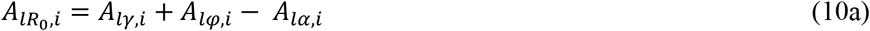

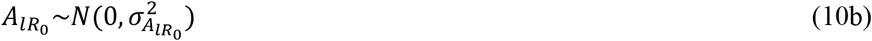

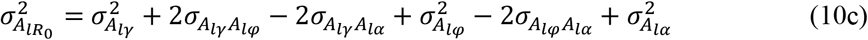

Hence, our model of the genotypic value for *R*_0_ is additive with Normally distributed effects on the log-scale. Thus the genotypic value for *R*_0_, as defined in Equations 8 and 9, follows a log-normal distribution,

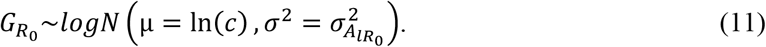

The genotypic value for *R*_0_ for the average individual, which has 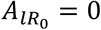, is equal to the contact rate, *c*. Hence, the genotypic value is defined here including its average, it is not expressed as a deviation from the mean. Moreover, we refer to 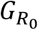 as a genotypic value, rather than a breeding value, because the 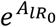 in Equation 9 is a non-linear function, so that 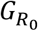 will show some non-additive genetic variance, even though 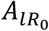 is additive.

The log-normal distribution of 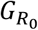 agrees with the infinitesimal model and the strictly positive values for *R*_0_ (Anacleto *et al*., 2015), and is also convenient because the mean and variance of 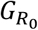 follow from the known properties of the log-normal distribution,

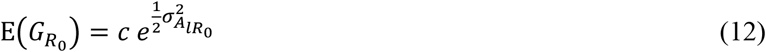

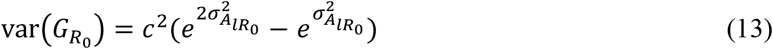

Equations 12 and 10c show that genetic (co)variation in susceptibility, infectivity and/or recovery, and thus in the breeding value for the logarithm of *R*_0_, leads to an increase in the mean genotypic value for *R*_0_. For example, for 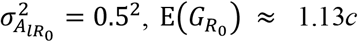. While this 13% increase in 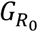 may suggest limited impact of heterogeneity, a 13% increase in *R*_0_ has a considerable impact on the endemic prevalence when *R*_0_ is close to one (Figure 1).

Equations 12 and 13 show that a log-normal distribution for susceptibility, infectivity and recovery results in a positive mean-variance relationship for 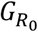. Figure 2 illustrates this relationship, for 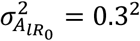 and genetic variation in susceptibility only. The *x*-axis shows the contact rate, which is equal to the genotypic value for *R*_0_ of the average individual. Hence, the *x*-axis reflects the level of *R*_0_. The small circle represents a population with a prevalence of ∼0.33, for which observed-scale heritability of binary infection status is ∼0.02 (Hulst *et al*., 2021). For that population, *R*_0_ is ∼1.5, and the genetic standard deviation in *R*_0_ is ∼0.48. Hence, despite the small observed-scale heritability, *R*_0_ has considerable genetic variation and some individuals will have a genotypic value smaller than 1, which agrees with the findings of Hulst *et al*. (2021). In the context of artificial selection against infectious diseases, the positive mean-variance relationship resulting from our model may be interpreted as conservative, because it implies a reduction of the genetic variance in *R*_0_ with continued selection for lower prevalence.

**Figure 2.**
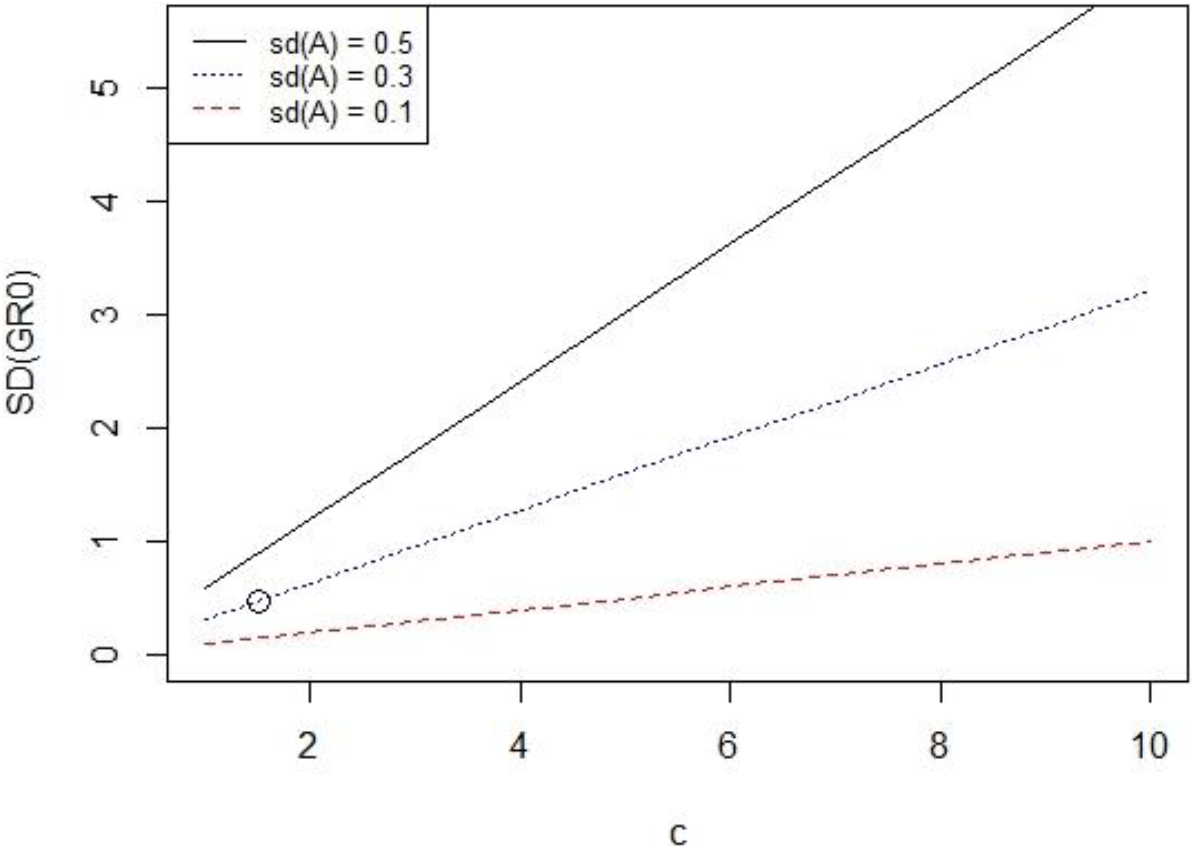
Genetic standard deviation in *R*_0_ as a function of the level of *R*_0_ (as approximated by the contact rate *c* here). From Equation 13, for three values of 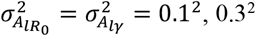 and 0.5^2^, and no variation in *φ* and α. The circle represents a population with a prevalence of ∼0.33, for which observed-scale heritability of binary infection status is ∼0.02 (Hulst *et al*., 2021).

In summary, this section has presented a genetic model for susceptibility, infectivity and recovery, leading to expressions for the genotypic value and genetic variance in *R*_0_ (Equations 8, 9 and 11). Note however, that we have not yet provided formal proof that the individual genotypic value for *R*_0_ indeed predicts the actual *R*_0_ (of the population so to say). In fact, the definition of 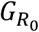 in Equation 8 is an educated guess based on the expression for *R*_0_ in a homogeneous population (Equation 4). In epidemiology, however, *R*_0_ is an emerging property of a population level process of the transmission of an infection, rather than an average of individual (genotypic) values. Thus it remains to be proven that the 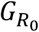 defined in Equation 8 indeed predicts the *R*_0_ of a genetically heterogeneous population. In the next two sections, therefore, we will focus on the population-level consequences of genetic heterogeneity, and investigate the impact of genetic variation on the level of *R*_0_ and on the endemic prevalence.

### 3. The impact of genetic heterogeneity on *R*_*0*_

*R*_0_ is a key parameter for infectious diseases, because infections can persist in a population if and only if *R*_0_ is greater than one (Kermack & McKendrik,1927; Diekmann *et al*. 1990). In other words, an endemic equilibrium can exist only when *R*_0_ is greater than one. Conversely, eradication of an infectious disease, either by vaccination or other measures such as genetic selection of the host population, requires that *R*_0_ is reduced to a value smaller than one. Here we address the consequences of genetic (co)variation in susceptibility, infectivity and recovery for the value of *R*_0_, and provide a proof that *R*_0_ is indeed the simple population average of the individual genotypic values for *R*_0_, as defined in Equations 8, 9 and 11. Note that *R*_0_ is strictly defined for the infection free state of the population (*i*.*e*., where the infected fraction is infinitesimally small). Hence, in this section we consider the infection free state, while the endemic equilibrium will be addressed in section 4.

Genetic (co)variation in susceptibility, infectivity and recovery has two consequences for *R*_0_. First, it increases the mean genotypic value for *R*_0_. This effect is trivial; it follows directly from Equations 12 and 10c and is not the main focus of this section. Second, as stated above, *R*_0_ is the average number of individuals that gets infected by a *typical* infected individual in an otherwise non-infected (large) population (Kermack and McKendrick, 1927; Diekmann *et al*., 1990). The expression for *R*_0_ given in Equation 4 ignores the “typical” term in the definition of *R*_0_, and is therefore an approximation in case of heterogeneity (Diekmann *et al*., 1990). The focus of this section is on the consequences of heterogeneity for the properties of the typical infected individual, and thus for *R*_0_.

The properties of the “typical infected individual” will depend on the magnitude and nature of the heterogeneity among the individuals in the population, because the susceptibility and recovery determine which animals are infected, while the infectivity of those individuals may differ from the population average. In contrast to the conclusion of Springbett *et al*. (2003), therefore, genetic heterogeneity can affect *R*_0_ (Diekmann *et al*. 1990, 2012). Suppose, for example, that individuals differ in both susceptibility and infectivity, and that susceptibility is positively correlated to infectivity. Because individuals with greater susceptibility are more likely to become infected, the typical infected individual will have an above-average susceptibility. Moreover, because of the positive correlation with infectivity, this will also translate into an above average infectivity of the typical infected individual, leading to higher *R*_0_. Hence, variation among individuals together with a positive (negative) correlation between susceptibility and infectivity results in an increase (decrease) in *R*_0_ (Diekmann *et al*. 1990). A similar argument holds for genetic covariation between recovery and infectivity. For this reason, *R*_0_ in general deviates from the right-hand side of Equation 4 obtained using the averages of *α, γ* and *φ*.

In Appendix 1, we derive the relationship between *R*_0_ and the genetic parameters for susceptibility, infectivity and recovery. The first step is the derivation of the lifetime infectivity of the typical infected individual. Lifetime infectivity is the total infectivity of an individual, aggregated over its average infectious period, and is the product of its infectivity per unit of time (*φ*, Equations 5 & 6) and the mean duration of its infectious period, 1/α_*i*_,

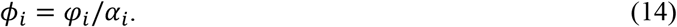

By introducing life time infectivity, we summarize the infectivity (per unit of time, *φ*) and the recovery of an individual into a single variable (*ϕ*), which simplifies the analysis. Appendix 1 shows that the average lifetime infectivity of the typical (*i*.*e*., average) infected individual equals

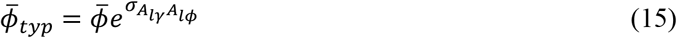

where 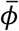 is the simple average of lifetime infectivity in the entire population, and 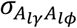 the covariance between the breeding values for the logarithms of susceptibility and lifetime infectivity. Thus Equation 15 is the average lifetime infectivity of infected individuals in the population at the infection free state, and can be interpreted as an average weighted for differences in susceptibility. It shows that the typical infected individual has an above (below) average life-time infectivity when the covariance between susceptibility and life-time infectivity is positive (negative), as argued verbally in the previous paragraph.

From the definition of *R*_0_ and Equations 4, 5 and 15, it follows that

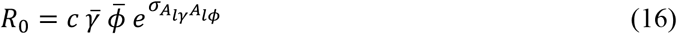

where 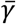 is the simple population average value of susceptibility. The last term of this expression shows that a positive covariance between susceptibility and life-time infectivity indeed increases *R*_0_. Note that this *R*_0_ is the reproduction number in the large infection free population, as it is normally defined in epidemiology. Hence the distribution of susceptibility in the susceptible individuals remains equal to the population distribution, in contrast to the distribution of infectivity in the infected individuals (See Appendix 1).

Equation 16 can be simplified by substituting the expression for 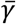 and 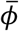, which follow from the log-normal distribution,

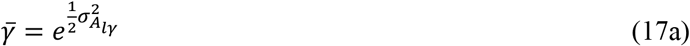

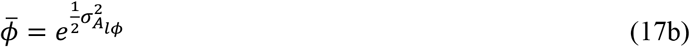

Substituting Equations 17a and b into Equation 16, and expressing the genetic variance of life-time infectivity in terms of infectivity per unit of time and recovery, reveals that *R*_0_ is equal to the simple population average of the individual genotypic value for *R*_0_ (see Appendix 1 and Equation 12),

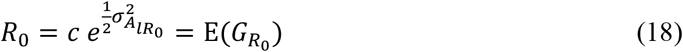

Thus, *R*_0_ depends on the variance in the breeding value for the logarithm of *R*_0_, but is still equal to the mean genotypic value for *R*_0_. In other words, a positive covariance between susceptibility and life-time infectivity indeed increases *R*_0_, but this effect is fully captured by the effect of the variance in the breeding values for the logarithm of *R*_0_ on the mean genotypic value for *R*_0_ (the 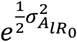 term in Equation 18). This result, therefore, provides formal proof that the genotypic value for *R*_0_, as defined in Equations 8 – 11, indeed represents the individual genetic value for *R*_0_.

Note that, while *R*_0_ is equal to the simple average genotypic value for *R*_0_, it still differs from the product of the simple averages of the rates when susceptibility, infectivity and/or recovery are correlated; 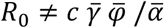. Moreover, *R*_0_ may also differ from the simple 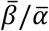 with heterogeneity. A numerical investigation of the 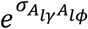 term in Equation 16 shows that a correlation between susceptibility and life time infectivity may change *R*_0_ by a maximum of about 25% for realistic levels of heterogeneity and log-normally distributed genetic effects. For example, for 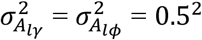, and a correlation 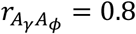 is 22% greater than 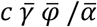. For values of *R*_0_ close to 1, this 22% may be the difference between absence of an infection *vs*. a significant endemic prevalence. Thus a correlation between susceptibility, infectivity and/or recovery may have a meaningful impact on *R*_0_.

In summary, this section has shown that heterogeneity and a positive correlation between susceptibly and life-time infectivity lead to an increase of *R*_0_, and thus increase the probability that an infectious disease persists in the population. However, when genotypic values for *R*_0_ follow a log-normal distribution, *R*_0_ is still equal to the simple average of those genotypic values.

### 4. The impact of genetic variation on the endemic prevalence

In this section we present an expression for the endemic prevalence in a population with genetic variation in susceptibility, infectivity and recovery, and also briefly investigate the quantitative effect of such variation for the endemic prevalence. Figure 1 and Equation 3 show the relationship between *R*_0_ and the endemic prevalence for a homogeneous population. With variation among individuals, however, more susceptible individuals are more likely to be in the infected state in the endemic equilibrium. For this reason, the mean susceptibility of the remaining non-infected individuals will be lower than the population average susceptibility. This in turn translates into an endemic prevalence lower than expected based on *R*_0_ (Equation 3; Diekmann *et al*. 2012; Springbett *et al*., 2003; Note, however, that the threshold value of *R*_0_ = 1 remains, so that endemic prevalence is zero if and only if *R*_0_ ≤ 1, and in that sense the *R*_0_ given in Equation 18 is exact). Similar arguments can be used to show that prevalence depends on the variation in the recovery rate, and on the covariation of infectivity, susceptibility and recovery. Thus Equation 3 is exact only in the absence of heterogeneity in these parameters.

The endemic prevalence in a heterogeneous population can be found by realizing that the prevalence must have reached an equilibrium value for each type of individual (Biemans *et al*., 2017; Aznar *et al*., 2018). Suppose, for example, that susceptibility, infectivity, and recovery would be governed by the same single bi-allelic locus in a diploid organism. Then, for the entire population to be in equilibrium, each of the three genotypic classes should be in equilibrium as well. In other words, the prevalence should have reached an equilibrium value within each genotypic class, but this value may differ among the three classes. Here we adapt this approach to continuous variation in polygenic traits.

In the endemic equilibrium, the number of susceptible individuals of each type, say *i*, should not change over time (apart from random fluctuation). Thus, for each type *i*, the number of newly infected susceptibles should be equal to the number of recovering infecteds,

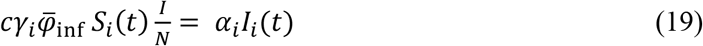

where *S*_*i*_ is the number of susceptible individuals of type *i, t* denotes time, *c* the contact rate, *γ*_*i*_ the susceptibility of type *i*, 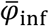 the mean infectivity of the infected individuals in the endemic equilibrium, *I* the total number of infected individuals in the endemic equilibrium, *N* total population size, α_*i*_ the recovery rate for type *i*, and *I*_*i*_ the number of infected individuals of type *i*. The left hand side in Equation 19 represents the decrease in the number of susceptibles due to transmission (infection), while the right hand side represents the increase of the number of susceptibles due to recovery of infected individuals. Our interest is in the solution of Equation 19 for *I*_*i*_ (or equivalently, for *S*_*i*_ = *N*_*i*_ – *I*_*i*_). Above, we used *i* to index individuals. Here we use *i* also to index types, since each individual will be genetically unique for polygenic traits, so that a type corresponds to an individual. Moreover, we treat *S*_*i*_ and *I*_*i*_ as non-integer because our interest is in their expectation. Note that the mean infectivity of the infected individuals in the endemic equilibrium 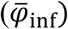 will differ from the simple population average of infectivity 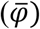 when infectivity is correlated to susceptibility and/or recovery.

Equation 19 can be solved for the endemic prevalence in type *i, P*_*i*_ = *I*_*i*_/*N*_*i*_, *N*_*i*_ denoting the total number of individuals of type *i* in the population,

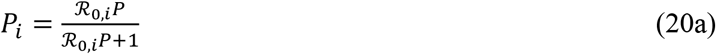

where *P* denotes the overall endemic prevalence in the population (Equation 1), and

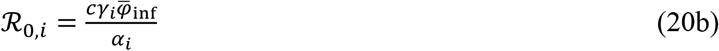

Equations 20a&b make no assumptions on the distribution of *γ, φ* and α, and are thus not restricted to log-normal distributions. Although Equation 20b is similar to Equation 8, note that ℛ_0,*i*_ differs from the genotypic value for *R*_0_ (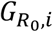; We use a symbol slightly different from *R* to highlight this difference). The ℛ_0,*i*_ is a function of the mean infectivity of the infected individuals in the endemic equilibrium 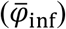, while 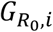 is a function of the infectivity of the individual itself (*φ*_*i*_). Our interest here is in the prevalence for an individual with susceptibility *γ*_*i*_ and recovery rate α_*i*_ in the endemic equilibrium, where *i* is exposed to the mean infectivity of the infected individuals. Hence the 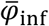 term in ℛ_0,*i*_. The 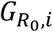, in contrast, defines the contribution of an individual’s genes to *R*_0_ (Equation 18), which is relevant for response to selection (section 7 below). Note that 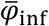 depends on the multivariate distribution of *γ, φ* and *α*.

To find the endemic prevalence, we need to solve Equations 20a and b for *P*. While we found an approximate analytical solution for the case without (correlated) genetic variation in infectivity, the resulting expression is very complex (not shown). We therefore used a numerical solution, which is easily obtained (see Appendix 2 for methods, and Supplementary Material 1 for an R-code). We validated the numerically obtained solution using full stochastic simulation of actual endemics, following standard methods in epidemiology. Results of these simulations confirmed the numerically obtained solutions (Appendix 3).

The solutions of Equations 20a and b show that variation in susceptibility and/or recovery reduces the endemic prevalence, compared to the simple prediction based on *R*_0_ (Equation 3). Hence, with variation in susceptibility and/or recovery, prevalence is always lower than predicted by Equation 3 (Figure 3; as expected with heterogeneity; Greenhalgh *et al*. 2000). Note that genetic variation in infectivity has no effect on the prevalence (beyond its trivial effect on the mean of 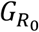, Equation 18), as long as infectivity is not correlated to susceptibility and/or recovery.

**Figure 3.**
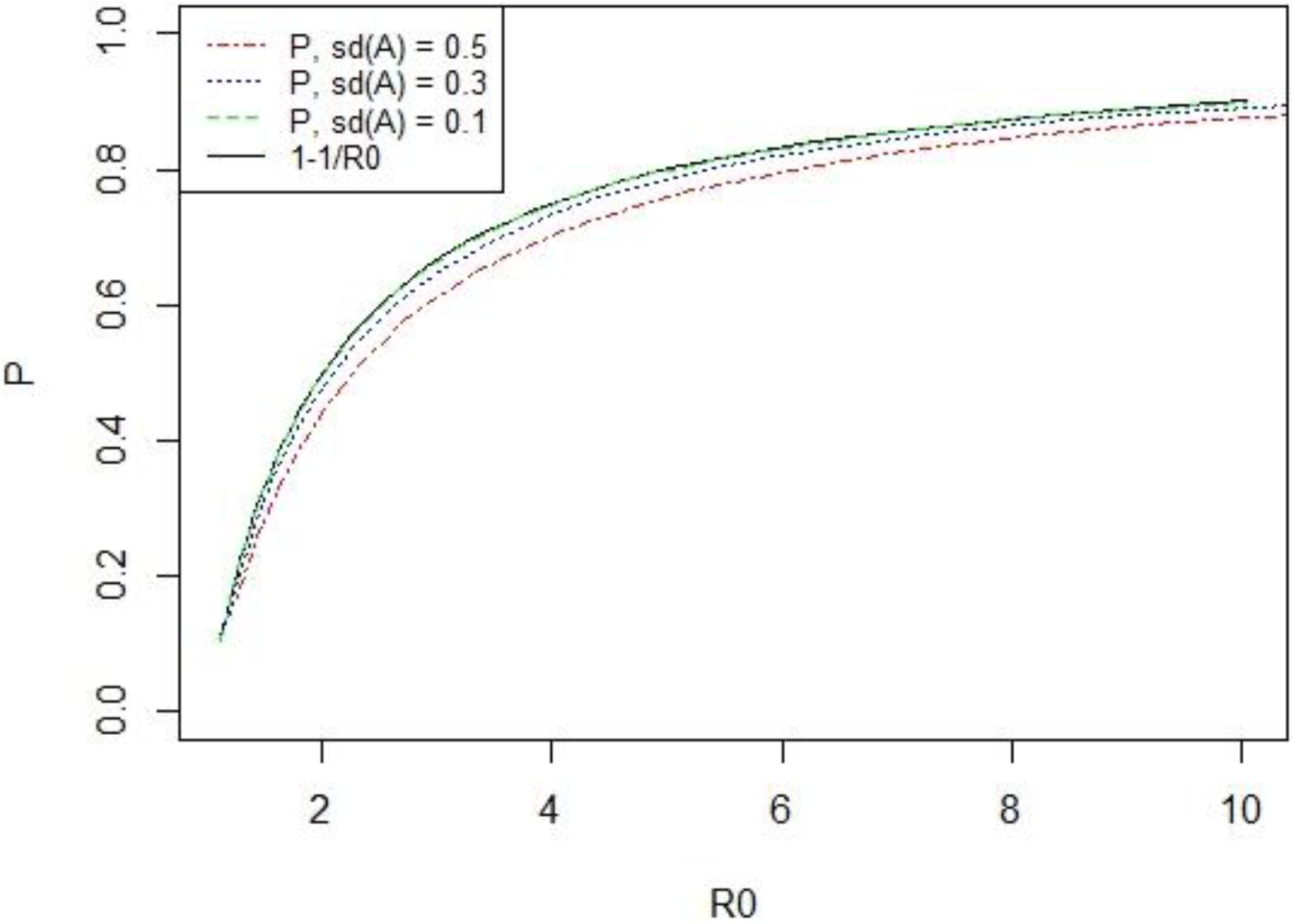
The impact of heterogeneity on the endemic prevalence. The solid line shows the prevalence predicted from Equation 3. The three other lines show the true prevalence (From numerically solving Equations 20a & b), for three levels of the additive genetic standard deviation in log susceptibility 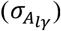, and no genetic variation in infectivity or recovery. The line for 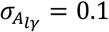 is almost identical to the solid line. Note that identical results would have been obtained with the same amount of heterogeneity in the recovery rate, or more generally in *A*_*lγ,i*_ − *A*_*l*α,*i*_, instead of in susceptibility.

Moreover, it follows from Equation 20 that the effects of genetic variation on the endemic prevalence are identical for susceptibility and recovery, since the ℛ_0,*i*_ of an individual depends on the difference between its breeding values for log-susceptibility and log-recovery, *A*_*lγ,i*_ − *A*_*l*α,*i*_,

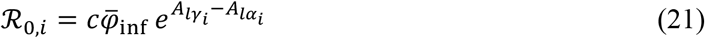

Hence, with a log-normal distribution of susceptibility and recovery, and in the absence of correlated variation in infectivity, the equilibrium prevalence depends only on (*R*_0_ and) the variance of this difference,

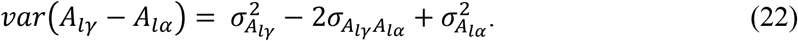

Figure 3 illustrates the impact of heterogeneity on the endemic prevalence for a limited number of scenarios with genetic variation in susceptibility only. For 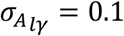, the effect of heterogeneity is imperceptible. For 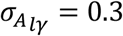, the true prevalence is up to 2 percent point lower than the value from Equation 3. For 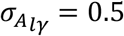, true prevalence is up to 6 percent point lower. This maximum difference occurs at a contact rate of two. Moreover, when *c* = 2 and there is no variation in infectivity, prevalence is always equal to 1 − 1/*c* = 0.5, irrespective of the genetic variation in susceptibility and recovery. (This is not visible in Figure 3, because the x-axis shows *R*_0_ rather than *c*). This occurs because the two opposing effects mentioned at the beginning of this paragraph exactly cancel each other. More detailed results can be found in Supplementary Material 2.

## 5. Genotypic value for individual binary infection status

In the previous two sections, we have considered the population-level effects of genetic heterogeneity. In the next two sections, we move to the individual level. This section focusses on the effects of an individual’s genes on its own infection status, while section 6 focusses on the effect of an individual’s genes on the prevalence in the population.

By definition, the genotypic value for binary disease status is the expected infection status of an individual given its genotype,

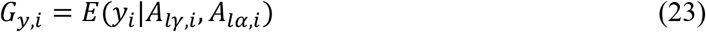

Thus, the *G*_*y*_ represents the direct genetic effect (DGE) on the own phenotypic value (including the mean, 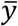, here; Note the distinction between subscript *y*, indicating individual binary infection status, and *γ*, indicating susceptibility). The genotypic value of an individual is not a function of its breeding value for log-infectivity, since an individual’s infectivity does not affect its own infection status. Hence, Equation 23 does not condition on *A*_*lφ,i*_.

In the previous section, we used Equations 20a and b to investigate the effect of heterogeneity on the endemic prevalence in the population. Equation 20a shows the expected prevalence of an individual of type *i*. However, since prevalence is simply the mean of binary infection status, Equation 20a may also be interpreted as the expected phenotypic value for infection status (*y* = 0,1) of an individual, given its genotype (specifically, the *γ*_*i*_ and *α*_*i*_ components of ℛ_0,*i*_). Hence, Equation 20a also represents the genotypic value for binary infection status,

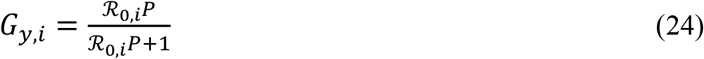

where ℛ_0 *i*_, follows from Equation 20b. The same result was found by Bijma (2020), but based on a different approach. Thus the *G*_*y*_ refers to the expected binary infection status of individual *i*, conditional on its genotype, in a population with prevalence *P*. Equations 21 and 24 imply that susceptibility and recovery are equally important for the infection status of an individual. For example, an individual with *A*_*lγ*_ = −0.1 has the same expected infection status as an individual with *A*_*lα*_ = +0.1.

Calculation of *G*_*y*_ from Equation 24 requires knowledge of the endemic prevalence *P*. In the previous section we used a numerical approach to find *P*, because our interest was in the effects of heterogeneity on *P*. In applied breeding, however, breeders may often have a reasonable idea of realistic values for the endemic prevalence, and a numerical solution may not be needed to find *G*_*y*_, or have little added value.

### Validation

We used stochastic simulation of endemics, following standard methods in epidemiology (Appendix 3), to validate Equation 24. Figures 4a-c show the mean observed infection status of individuals as a function of their genotypic value *G*_*y*_. For all three panels in Figure 4, regression coefficients were very close to 1, showing that *G*_*y*_ is an unbiased linear predictor of individual infection status.

**Figure 4.**
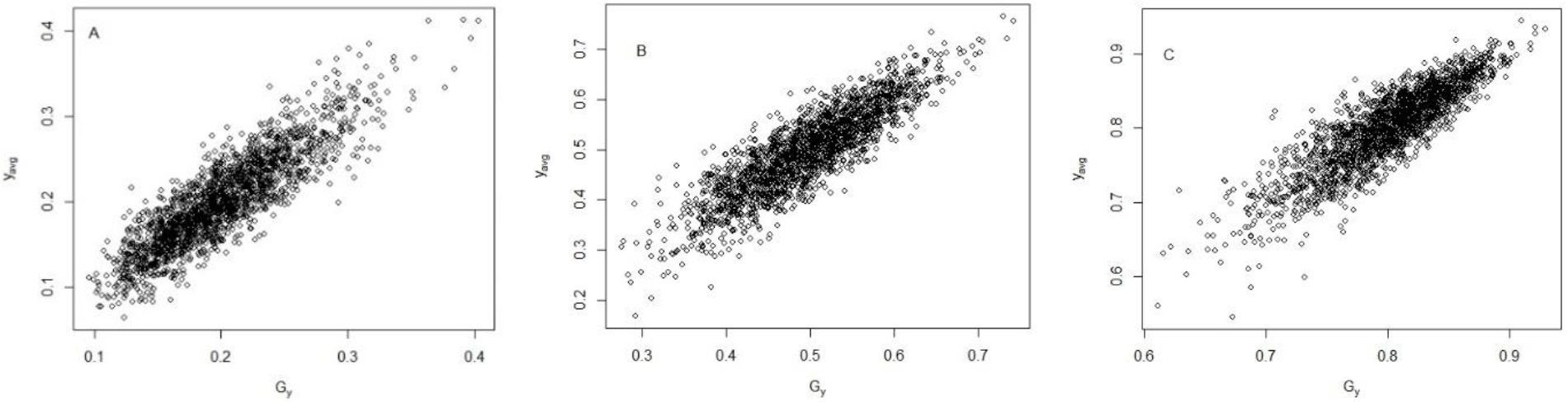
Validation of the genotypic value for individual binary infection status. Panels show a scatter plot of the mean observed infection status of individuals (y-axis) as a function of their genotypic value for infection status (*G*_*y*_, x-axis, Equation 24). For genetic variation in susceptibility only, with 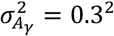, *N* = 2000 individuals, and a total of 300,000 events (sum of recoveries and infections). Panel A: 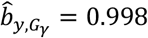. Panel B: P = 0.5, 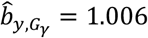. Panel C: P = 0.8, 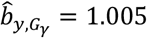

We numerically investigated the relative amount of non-additive genetic variance in *G*_*y*_. Results (not shown) revealed only little non-additive genetic variance. For example, for *P* = 0.2 and 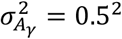, more than 96% of the genotypic variance in *y* was additive. Thus the breeding value for own infection status is very similar to the genotypic value,

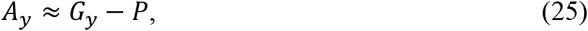

where the “−*P*” term simply reflects subtraction of the average,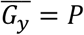, so that the mean breeding value is zero by definition. We defer further investigation of the breeding value and the additive genetic variance for individual infection status to the next section, to facilitate comparison with the corresponding measures for prevalence.

## 6. Breeding value and heritable variation for the endemic prevalence

The previous section focussed on the genetic effects of individuals on their own infection status. In this section we will consider the genetic effects of individuals on the endemic prevalence in the population. In other words, the previous section focussed on the contribution of genetic effects to the variation in infection status among individuals, while this section considers the genetic effects that are relevant for response to selection. We will present expressions for the genotypic value, breeding value and additive genetic variance for the endemic prevalence. The genotypic value will reflect the full genetic effect of an individual on the endemic prevalence in the population, while the breeding value reflects the additive component thereof. The last part of this section contains a comparison of the breeding value for endemic prevalence and that for individual infection status.

The relationship between *R*_0_ and the endemic prevalence (Equation 3) suggests we can translate the individual genotypic value for *R*_0_ (Equations 8 and 9) to the scale of prevalence, by defining an individual genotypic value for prevalence as

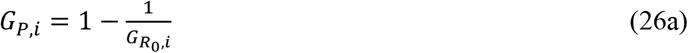

Because this definition is based on Equation 3, it ignores the of effect heterogeneity on the relationship between *P* and *R*_0_ (Figure 3). We will investigate the relevance of this approximation numerically in section 7, which focusses on response to selection. Substituting Equation 9 into 26a yields an expression for the genotypic value of an individual for the endemic prevalence, in terms of its breeding value for the logarithm of *R*_0_,

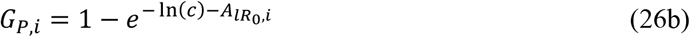

Because the term in the exponent is Normally distributed, 1 − *G*_*P*_ follows a log-normal distribution,

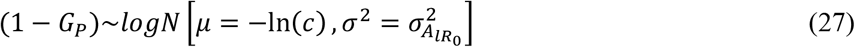

The mean and variance of the genotypic values for prevalence, therefore, follow from the properties of the log-normal distribution,

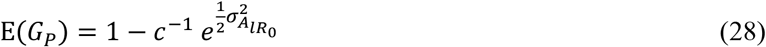

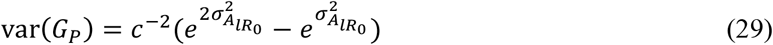

To enhance the interpretation of Equation 29, we can express it as a function of *R*_0_ or of the endemic prevalence. Substituting Equation 18 into Equation 29 yields an expression for the genetic variance in prevalence as a function of *R*_0_,

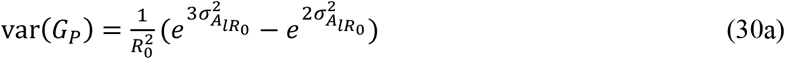

Given the definition of *G*_*P*_ in Equation 26a, Equation 30a is exact. Next, substituting Equation 3 into Equation 30a yields an expression for var(*G*_*P*_) as a function of the endemic prevalence,

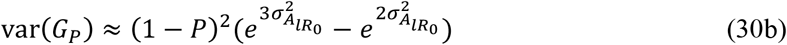

Equation 30b is approximate, because the relationship between *P* and *R*_0_ given in Equation 3 is approximate with heterogeneity. Equations 30a and b show how the genotypic variance for endemic prevalence depends on the level of *R*_0_ or equivalently, on the level of the endemic prevalence. Hence, in contrast to ordinary additive genetic traits, the genetic variance for endemic prevalence is a function of the level of the endemic prevalence (Equation 30b).

Figures 5a and b illustrate that the standard deviation in genetic values for endemic prevalence is considerably larger at lower *R*_0_, or equivalently, at lower prevalence. Hence, even though the genetic variance in *R*_0_ decreases with the level of *R*_0_ (Figure 2), the genetic variance for prevalence increases strongly when *R*_0_ decreases. This result originates from the increasing slope of the relationship between prevalence and *R*_0_ when *R*_0_ decreases (Figure 1). In other words, an equal change in *R*_0_ has much greater impact on the endemic prevalence at low *R*_0_ than at high *R*_0_, which is well-known in epidemiology (*e*.*g*., Metz, 1978; Bolker and Grenfell, 1996). Hence, for a constant variance in the breeding value for the logarithm of *R*_0_, the genetic variance for endemic prevalence is much greater at lower prevalence. Moreover, genetic selection for lower prevalence will lead to an increase in the genetic variance for prevalence.

**Figure 5.**
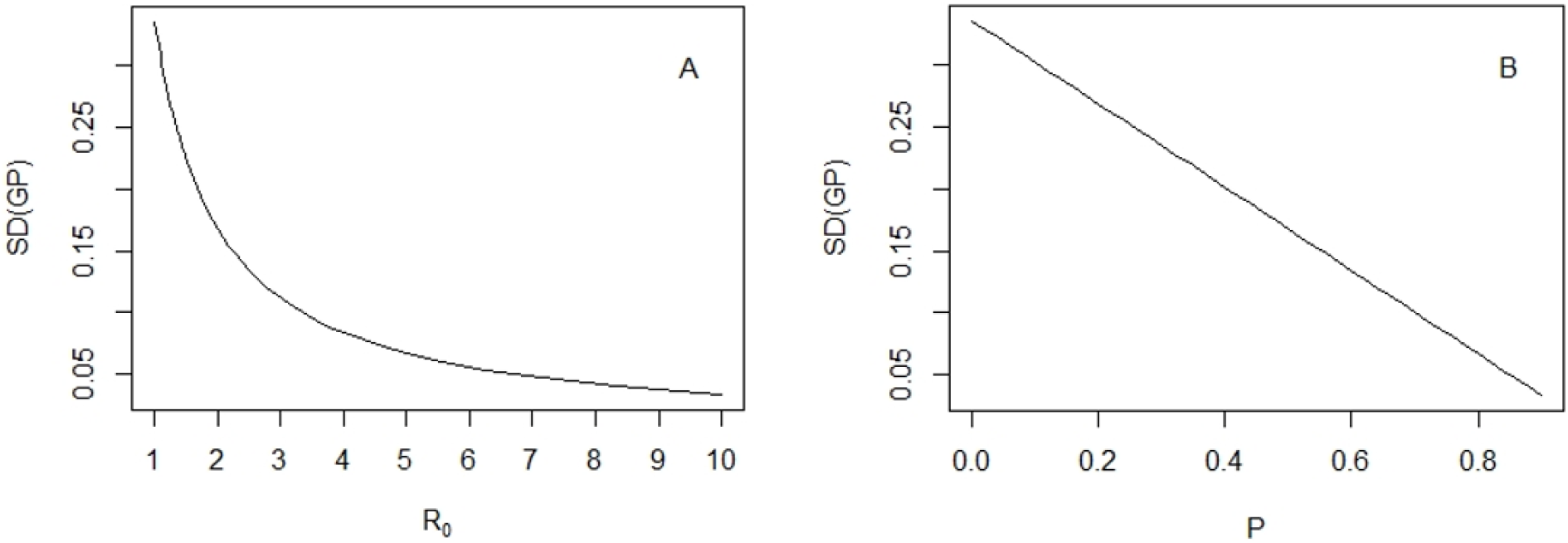
Genetic standard deviation for endemic prevalence as a function of *R*_0_ (panel A), and as a function of the level of the endemic prevalence (panel B). From Equations 30a and b. For 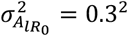. In panel A, *x*-axis values below *R*_0_ = 1 are omitted, because equilibrium prevalence is zero (the infection is not present) and Equation 30a does not apply.

Figure 6 shows some examples of the distribution of the genotypic value for endemic prevalence, for different values of *R*_0_ and the corresponding endemic prevalence. For the scenarios in Figure 6, the observed-scale heritability of individual infection status does not exceed 0.022 (see Figure 7 below). The panels illustrate that the genotypic standard deviation for endemic prevalence is relatively large, particularly when prevalence is small. For example, for *R*_0_ = 1.67 (*P ≈* 0.4; Panel B), the standard deviation in genotypic values for prevalence is around 0.19 (See also Figure 5), and values between ∼0 and ∼0.7 are quite probable. Hence, despite the low observed-scale heritability of individual infection status, the probable values of *G*_*P*_ span as much as 70% of the full 0-1 range of endemic prevalence.

**Figure 6.**
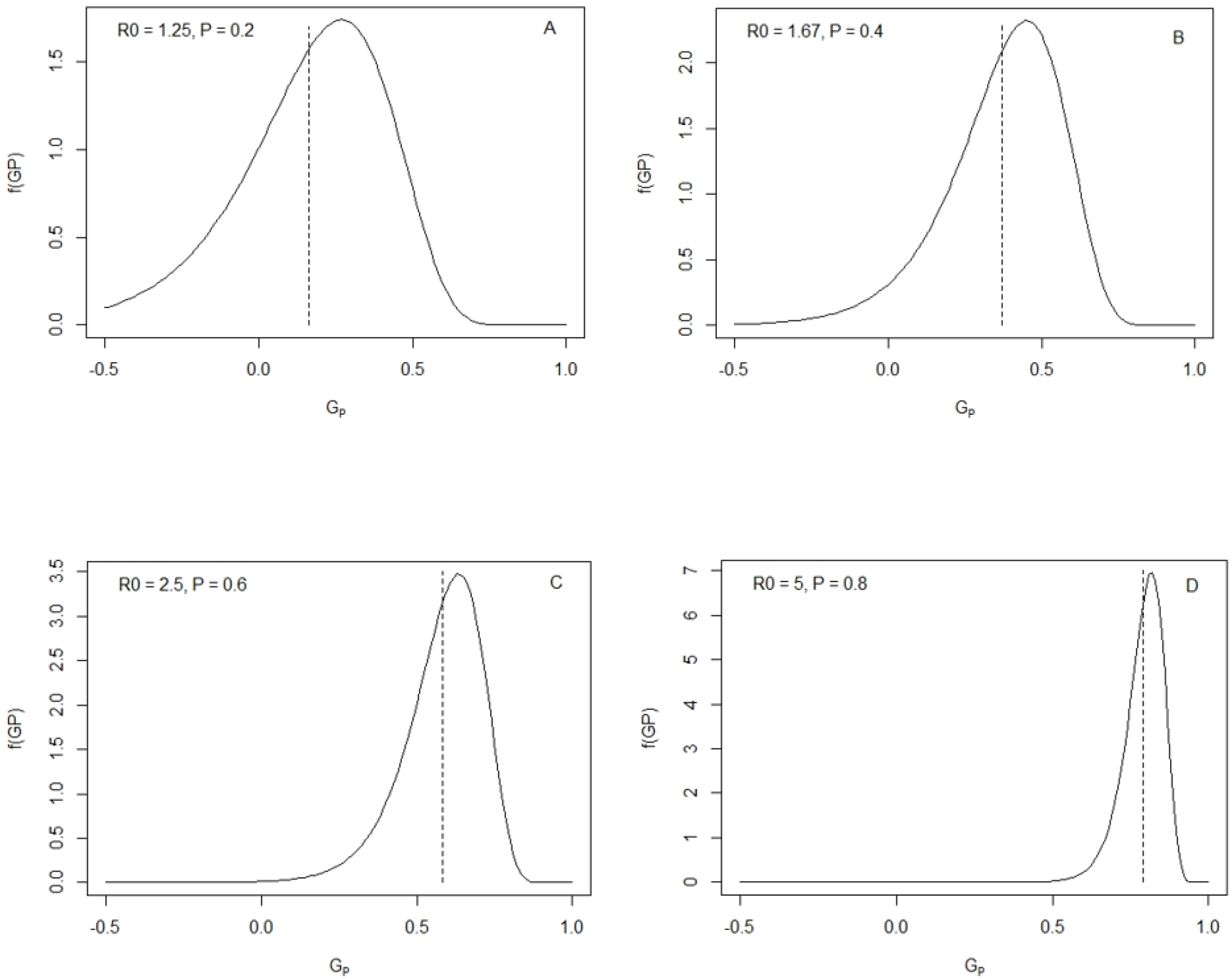
Distribution of individual genotypic values for prevalence (*G*_*P*_), for different values of *R*_0_, or equivalently, different (approximate) values of the endemic prevalence. The distribution is given by 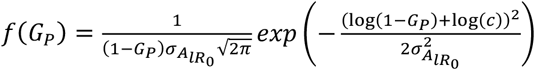, with domain *G*_*P*_= (-∞, 1). For 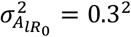. The dashed vertical line shows the mean of *G*_*P*_. Note that *G*_*P*_ can take negative values while prevalence cannot. This is because *G*_*P*_ reflects the genetic effect of an individual on the prevalence of the population, not the expected value of its own infection status. Thus, negative values for *G*_*P*_ are possible, as long as *P* is positive. Note that *P* is very close to the average of the distributions shown.

**Figure 7.**
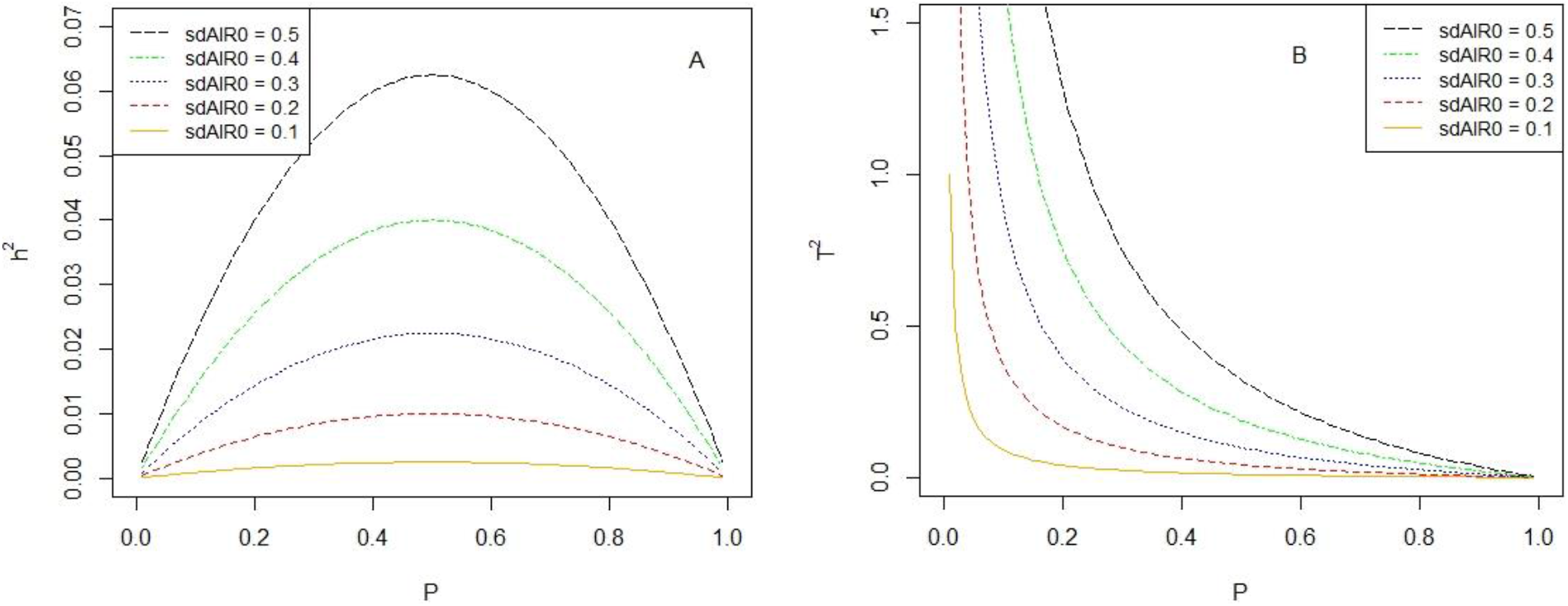
Panel A: Observed-scale heritability 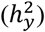 of individual binary infection status (*y* = 0,1) as a function of the endemic prevalence, for different additive genetic standard deviations in the logarithm of *R*_0_ (SDAlR0). From Equation 36. Panel B: Ratio of additive genetic variance for prevalence and phenotypic variance in infection status 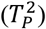, as a function of the endemic prevalence. From Equations 37 and 32a. In both panels, there is no genetic variation in infectivity.

### Breeding value and additive genetic variance for prevalence

The genotypic value for prevalence is not identical to the additive genetic value (*i*.*e*., breeding value) for prevalence, because the exponential function in Equation 26b is non-linear, so that *G*_*P*_ contains a non-additive component. Appendix 4 shows that the linear regression coefficient of *G*_*P*_ on 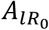 is equal to 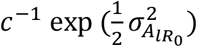. Substituting 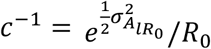 (from Equation 18), shows that the breeding value for prevalence is given by

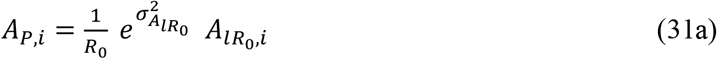

Given the definition of *G*_*P*_ in Equation 26a, this result is exact. With limited heterogeneity, this result is approximately equal to

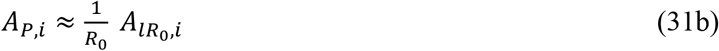

Thus the breeding value for prevalence is proportional to the reciprocal of *R*_0_. The additive genetic variance for prevalence equals

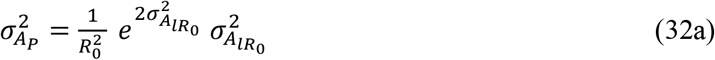

and, with limited heterogeneity,

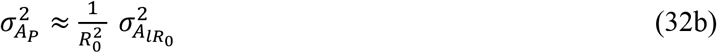

or, expressed as a function of endemic prevalence,

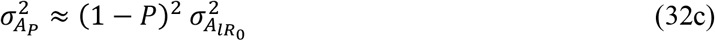

Equation 32c shows that the additive genetic variance in endemic prevalence increases strongly when prevalence decreases, similar to the relationship between the genotypic variance and endemic prevalence (Figure 5). This result suggests that response of endemic prevalence to selection will be greater at lower levels of the prevalence, which we will further investigate in Section 7 below.

The relative amount of non-additive genetic variance in the endemic prevalence is determined by the magnitude of 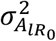 (Appendix 4). For realistic values of 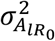, the vast majority of the genotypic variance in prevalence is additive. For example, for 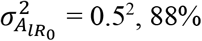, 88% of the variance in *G*_*P*_ is additive. Hence, the distinction between the breeding value for prevalence (*A*_*P*_) and the genotypic value for prevalence (*G*_*P*_) seems of minor importance, and results in Figures 5 and 6 will closely resemble those for the additive genetic effects.

### Breeding value and heritability for infection status *vs*. prevalence

Appendix 5 shows that, in the absence of genetic variation in infectivity, the breeding value for endemic prevalence is approximately a factor 1/*P* greater than the breeding value for individual infection status,

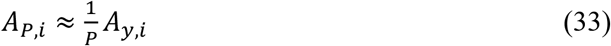

Note, in contrast to genotypic values, breeding values are expressed as a deviation from their mean here. The *A*_*y*_ is the ordinary observed-scale breeding value for binary infection status that breeders are familiar with.

Equation 33 implies that the impact of an individual’s genes on the response of the endemic prevalence to selection is considerably larger than their impact on the infection status of the individual itself, particularly when the endemic prevalence is small. Consider, for example, an individual with *A*_*y,i*_ = −0.02 in a population with an endemic prevalence of *P* = 20%. The expected infection status of this individual in the current population equals 0.20 − 0.02 = 0.18. Hence, on average, this individual will be infected 18% of the time. However, its breeding value for prevalence equals *A*_*P,i*_ = −0.02/0.2 = −0.10. Hence, if we select individuals with *A*_*y,i*_ = −0.02 as parents of the next generation, then the endemic prevalence will go down to 0.20 – 0.10 = 0.10. In other words, the response to selection will be fivefold greater than suggested by the ordinary breeding value for individual infection status (since 1/*P* = 1/0.2 = 5). We will numerically validate this theoretical result in Section 7.

The relationship between the breeding value for prevalence and the breeding value for own infection status shown in Equation 33 suggests a relatively simple expressions for *A*_*y*_. Such a simple expression would be convenient, because the alternative is to calculate *A*_*y*_ from Equation 25, which requires solving Equations 20a and b numerically. On combining Equations 31a and 33, and assuming limited heterogeneity, so that 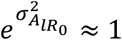 and 1/*R*_0_ ≈ (1 − *P*), the breeding value for individual infection status becomes

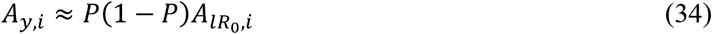

We used stochastic simulation to validate this expression and investigate its precision. Results show that Equation 34 closely matches the regression of individual binary infection status on the breeding value for the logarithm of *R*_0_ for realistic levels of heterogeneity (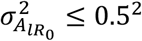; Supplementary Material 3; the good fit results from compensating errors due to the approximations). Hence, Equation 34 is sufficiently precise for practical purposes. Note that, since infectivity does not affect the infection status of an individual itself, a potential component due to infectivity has to be left out of the 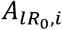 term when calculating Equation 34. In other words, in Equation 34 the 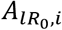 should include only the breeding values for the logarithm of susceptibility and recovery (see Equation 10a).

It follows from Equation 34 that the additive genetic variance in individual binary infection status equals

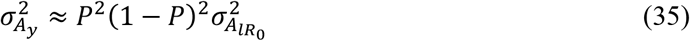

Next, the observed-scale heritability of binary infection status follows from dividing Equation 35 by the phenotypic variance of binary infection status, 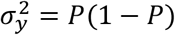, giving

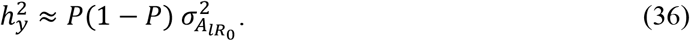

Hence, the observed-scale heritability for binary infection status has a maximum at a prevalence of 0.5, and goes to zero at a prevalence of zero or one, just like the heritability of binary phenotypes for non-communicable polygenic traits (Robertson, 1950; Figure 7A; assuming the infinitesimal model at the level of the logarithm of *R*_0_, so that 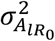 is constant).

The ratio of additive genetic variance for prevalence over phenotypic variance in binary infection status is given by,

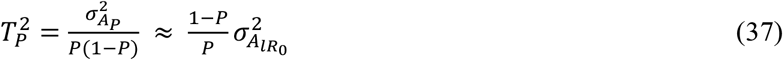

with 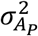 taken from Equation 32a or c. The 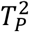 is an analogy of heritability, but the numerator represents the additive genetic variance relevant for response to selection in endemic prevalence, rather than for individual binary infection status. The 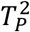, therefore, reflects the genetic variance that can be used for response to selection, whereas 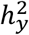 reflects the contribution of additive genetic effects to phenotypic variance (Bijma *et al*., 2007; Bijma 2011).

Figures 7 A & B show a comparison of 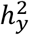 and 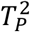 for a population without genetic variation in infectivity, with genetic variances in the logarithm of *R*_0_ ranging from 0.1^2^ through 0.5^2^. In Figure 7A, the maximum value of 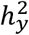 equals 0.0625, for *P* = 0.5 and 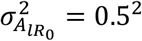. Given that genetic variances greater than 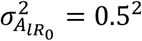 are very large (as argued above), observed-scale heritabilities of binary infection status greater than ∼0.06 are unlikely for endemic infectious diseases. The heritabilities in Figure 7A agree with the findings of Hulst *et al*. (2021), who used stochastic simulation of actual endemics and analysis of the resulting binary infection status data with a linear animal model. Figure 7B shows that 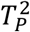 increases strongly when prevalence goes down. Figure 7 illustrates that 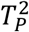 and 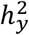 differ by a factor of approximately *P*^2^, so that the additive genetic variance in prevalence is (much) greater than the additive genetic variance in individual infection status, and may even exceed the phenotypic variance at low values of the endemic prevalence (*i*.*e*., 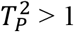).

In conclusion, in this section we have presented expressions for the breeding value for prevalence (Equation 31) and for individual infection status (Equation 34), and for the corresponding genetic variances. With realistic levels of heterogeneity, the breeding value for prevalence is a factor 1/*P* greater than the breeding value for individual infection status. This result suggests that response to selection should be considerably greater than expected based on ordinary heritability of individual infection status. We will test this hypothesis in the next section.

## 7. Response to selection

The higher genetic variance for prevalence at lower values of the prevalence (Equations 30 & 32, Figure 5b & 6) suggests that response to selection should increase when the prevalence decreases. To validate and illustrate this hypothesis, we stochastically simulated an endemic infectious disease in a large population undergoing mass selection for individual infection status. Simulations were based on standard methods in epidemiology, not making use of the above theory (Appendix 6).

Figure 8A shows the observed prevalence (*i*.*e*., the mean binary infection status in each generation), the mean breeding value for prevalence and the mean breeding value for binary infection status, for ∼70 generations of selection. Response in prevalence increases strongly when prevalence decreases, and the infection disappears in the final generation. There is excellent agreement between the observed prevalence and the breeding value for prevalence, showing that the change in Ā_*P*_ indeed predicts the change in prevalence. In contrast, the response in prevalence deviates substantially from the response in the breeding value for individual infection status (Ā_*y*_), particularly at lower values of the prevalence. Hence, while the breeding value for infection status correctly predicts the average individual infection status within a generation (Figure 4), the change in Ā_*y*_ considerably underestimates the response to selection. Furthermore, given the weak selection and the low value of the observe-scale heritability of binary infection status, which did not exceed 0.022 in Figure 8A, response to selection in prevalence is remarkably large, unless prevalence is high. This result agrees with findings of Hulst *et al*. (2021).

**Figure 8.**
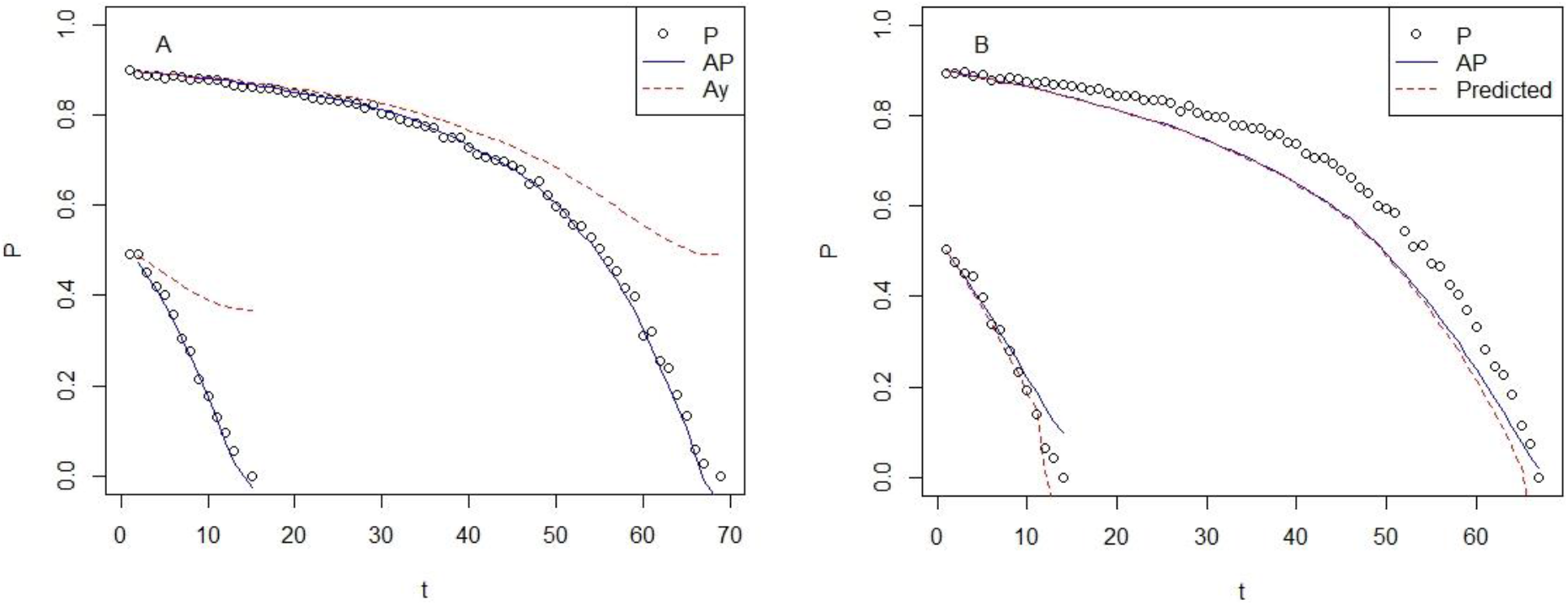
Response to selection in prevalence for 70 generations of mass selection of the host population. For two populations, one starting at a prevalence of ∼90% (*c* = 10), the other starting at a prevalence of ∼50% (*c* = 2). Each generation, the 50% individuals with the lowest average infection status were selected as parents of the next generation. With genetic variation in susceptibility only, and 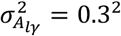. For a population of *N* = 4,000 individuals, a total of 15,000 events (sum of infections and recoveries) per generation, consisting of a burn-in of 10,000 events and 5,000 recorded events. Hence, selection is based on 1.25 events per individual on average, indicating a limited amount of phenotypic data. Observed-scale heritability for binary infection status in any generation can be read from Figure 7A using an *x*-axis value corresponding to the prevalence in that generation. *Panel A:* Observed prevalence (circles), breeding value for prevalence (Ā_*P*_, solid blue line) and breeding value for individual infection status (Ā_*y*_, dashed red line). Results for breeding values are the cumulative change in breeding value in each generation plus the initial prevalence. Breeding values were taken from Equation 31a for *A*_*P*_ and from Equation 34 for *A*_*P*_. *Panel B*: Predicted (lines) versus observed (circles) prevalence. Prevalence was predicted from Equation 38 (blue solid line) or Equation 39 (red dashed line).

We also briefly investigated the prediction of response to mass selection with a very simple expression assuming linearity and with genetic variation in susceptibility only. We based our predictions on breeding values for binary infection status, because these are typically estimated by breeders. Because the breeding value for prevalence and the breeding value for individual infection status differ by a factor 1/*P* when there is no genetic variation in infectivity (Equation 33), we simply upscaled the response to selection predicted for individual infection status from the breeder’s equation (Walsh and Lynch, 2018) by this factor, giving

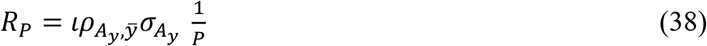

where *l* is the intensity of selection, defined as the standardized selection differential in mean individual infection status, 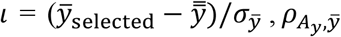 is the accuracy of selection for individual infection status, which is the correlation between the selection criterion 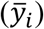 and the true breeding value for individual infection status, and *P* is the prevalence in the generation of the selection candidates. Hence, the numerator of Equation 38 represents the predicted response for individual binary infection status, which is multiplied by a factor 1/*P* to find response in prevalence. To implement Equation 38, we calculated the 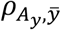 as the observed correlation between the true breeding values for binary infection status (*A*_*y*_; Eqn. 34) and the selection criterion 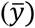 in the candidates for selection. Hence, we did not attempt to predict the accuracy of selection.

Figure 8B shows a comparison of observed and predicted prevalence. Above a prevalence of ∼0.5, response predicted from Equation 38 is somewhat larger than observed response, while the reverse is true below a prevalence of ∼0.5 (Note, response to selection in a generation is reflected by the slope of the figure). Nevertheless, agreement between observed and predicted response is remarkably good given the very unrealistic assumption of linearity in Equation 38 (*i*.*e*., bivariate normality of *A*_*y*_ and 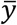). Because selection was based on mean individual infection status recorded over a period lasting on average only 1.25 events per individual (see legend Figure 8), many values were either 0 or 1, implying strong deviations from normality.

When the prevalence was smaller than 0.5, response to selection was quite large. Hence, there was a meaningful difference in prevalence between parent and offspring generations. Because the *P* in Equation 38 refers to the prevalence in the parent generation, while response is realized in the offspring generation, Equation 38 resulted in underprediction of response to selection when response was large. This underprediction disappeared when using prevalence in the offspring generation in the 1/*P* term in Equation 38. However, because prevalence in the offspring generation is initially unknown, as it depends on the response to selection, this prediction required solving the expression 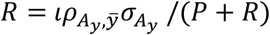, yielding

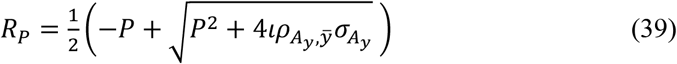

For a prevalence smaller than ∼0.5, predictions from Equation 39 were very close to the observed response in prevalence (Figure 8B; for *P* > 0.5, results of Equations 38 and 39 are almost identical).

In conclusion, results in this section show that response to selection in the prevalence of endemic infectious diseases is a factor 1/*P* greater than suggested by the ordinary breeding values for individual binary infection status. Thus breeders can predict response to selection by upscaling the selection differential in the usual estimated breeding values for binary infection status by a factor 1/*P*.

## 8. Direct and indirect genetic variance for endemic prevalence

In this section, we partition the total additive genetic variance for endemic prevalence into direct and indirect genetic components. This partitioning is relevant, because IGE respond fundamentally different to selection than DGE (Griffing 1967, 1977; Wright 1985; Moore *et al*., 1997; Muir 2005; Bijma 2010, 2011; see Discussion). We can partition the total breeding value for prevalence into a direct and an indirect component,

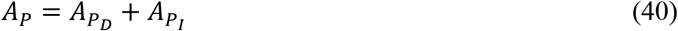

Analogously, we can partition the full additive genetic variance in prevalence into components due to direct genetic variance, indirect genetic variance and a covariance,

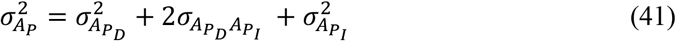

In the absence of genetic variation in infectivity, the breeding value for own infection status is a fraction *P* of the breeding value for prevalence (Equation 33). Hence, a fraction *P* of the additive genetic effects of susceptibility and recovery on prevalence affects the infection status of the individual itself and thus represents a direct effect, while the remaining fraction (1 − *P*) represents an indirect effect. For infectivity, the entire genetic effect is indirect, because an individual’s infectivity does not affect its own infection status. It follows from Equations 31a that

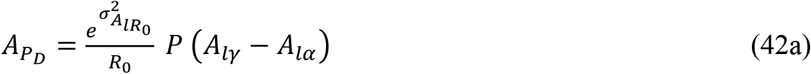

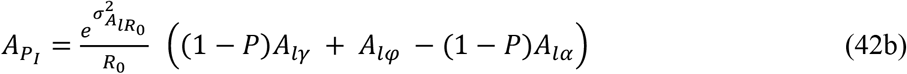

Note that Equation 42a represents the breeding value for individual infection status (*A*_*y*_), but the current expression emphasizes the partitioning of *A*_*P*_ into direct and indirect effects. The direct and indirect genetic (co)variances are given by

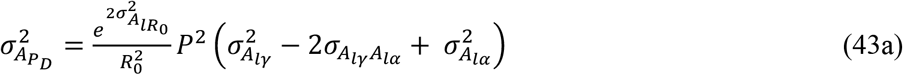

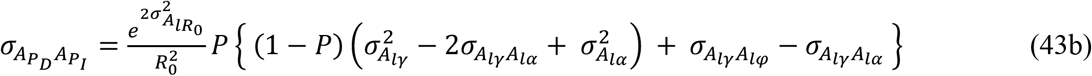

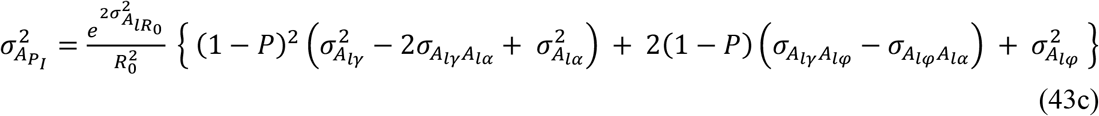

Figure 9 shows the total additive genetic variance for the endemic prevalence and the fractions due to DGE, IGE and their covariance, for a scenario with equal genetic variances in susceptibility, infectivity and recovery and covariances equal to zero. For an endemic prevalence smaller than 0.5, IGE contribute the majority of the genetic variance. For example, for an endemic prevalence of 0.3, the total additive genetic variance consists of 6% direct genetic variance, 66% indirect genetic variance and 28% direct-indirect genetic covariance. These results imply that IGE dominate the heritable variation and response to selection for the endemic prevalence of infectious diseases, unless prevalence is high.

**Figure 9.**
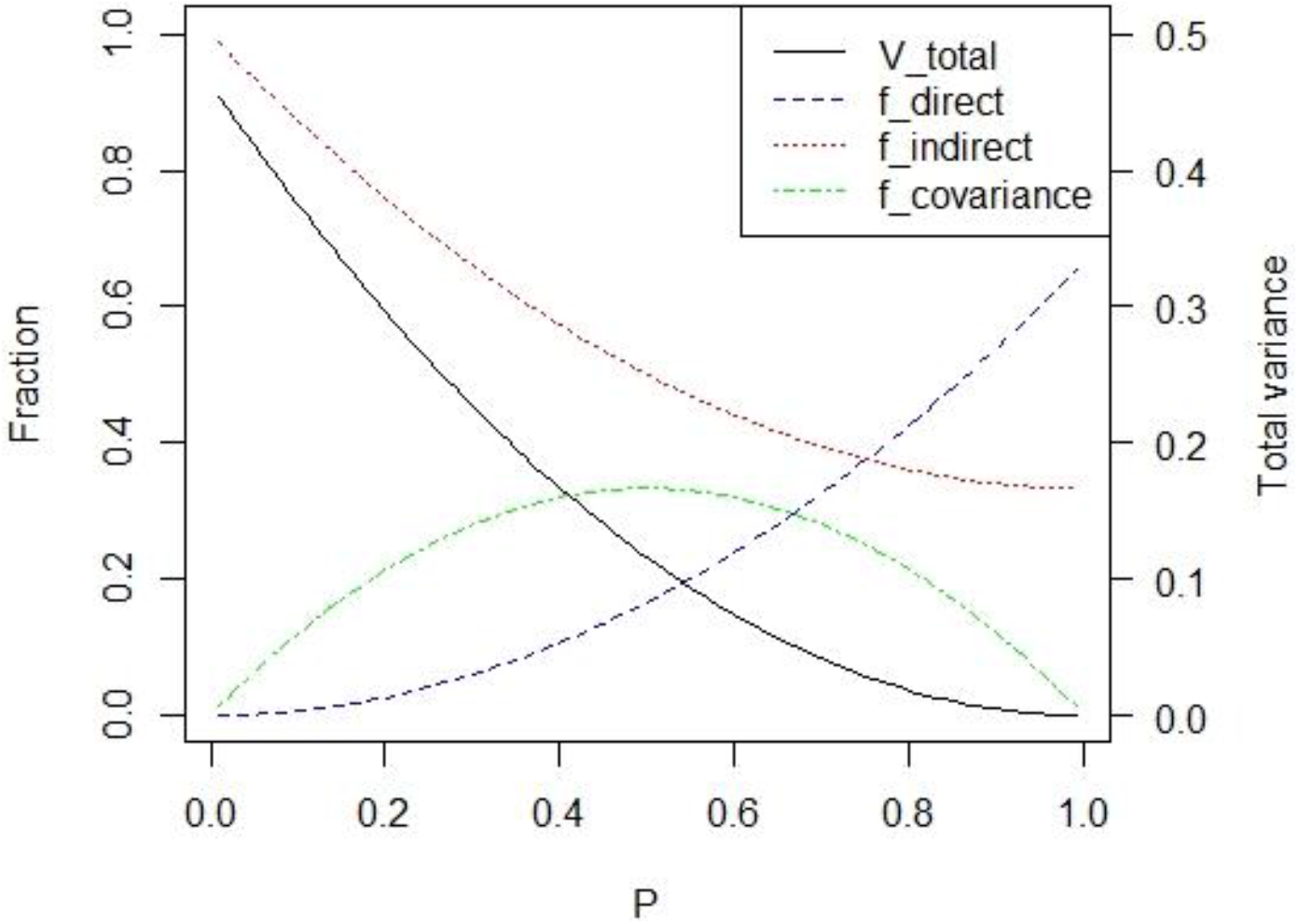
Total additive genetic variance in endemic prevalence (*V*_total_; secondary y-axis) and the relative contributions of DGE, IGE and their covariance (*f*_direct_, *f*_indirect_, and *f*_covariance_; primary y-axis). For constant values of 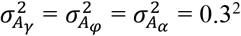 and covariances equal to zero. Results are obtained from Equations 32b, 3, and 43a-c.

## Discussion

We have presented a quantitative genetic theory for endemic infectious diseases, with a focus on the genetic factors that determine the endemic prevalence. We defined an additive model for the logarithm of individual susceptibility, infectivity and rate of recovery, which results in normally distributed breeding values for the logarithm of *R*_0_. Next, we investigated the impact of genetic heterogeneity on the population level, for both *R*_0_ and the endemic prevalence. Results show that, despite heterogeneity, *R*_0_ remains equal to the mean individual genotypic value for *R*_0_. Subsequently, we considered genetic effects of individuals on their own infection status and on the endemic prevalence in the population. Building on the breeding value for the logarithm of *R*_0_, we showed that genotypic values and genetic parameters for the prevalence follow from the known properties of the log-normal distribution. In the absence of genetic variation in infectivity, genetic effects for the endemic prevalence are a factor 1/prevalence greater than the ordinary genetic values for individual binary infection status. Hence, even though prevalence is the simple average of individual binary infection status, breeding values for prevalence show much more variation than those for individual infection status. These results imply that the genetic variance that determines the potential response of the endemic prevalence to selection is largely due to IGE, and thus hidden to classical genetic analysis and selection. For susceptibility and recovery, a fraction 1-*P* of the full genetic effect on endemic prevalence is due to IGE, whereas the effect of infectivity is entirely due to IGE. Hence, the genetic variance that determines the potential response of the endemic prevalence to selection must be much greater than expected based on classical quantitative genetic theory, particularly at low levels of the prevalence (Figure 7). We evaluated this implication using stochastic simulation of endemics following standard methods in epidemiology, where parents of the next generation were selected based on their own infection status (mass selection). The results of these simulations show that response to selection in the observed prevalence and in the breeding value for prevalence increases strongly when prevalence decreases, and closely matches our predictions, which supports the theoretical findings presented here.

### Model assumptions

Following Anacleto *et al*. (2015, 2019), Biemans *et al*. (2019) and Pooley *et al*. (2020), we assumed a linear additive model with normally distributed effects for the logarithm of susceptibility, infectivity and recovery, leading to a normal distribution of the additive genetic values for the logarithm of *R*_0_ (Equation 10). For complex traits, it is common to assume normally distributed genetic effects, based on the central limit theorem (Fisher 1918). Because susceptibility, infectivity and recovery act multiplicatively in the expression for *R*_0_, and because *R*_0_ is non-negative, we specified a normal distribution for its logarithm. This resulted in an additive model on the log scale, which fits with the infinitesimal model, and also translates the [0, ∞) domain of *R*_0_ to the (−∞, +∞) domain of the normal distribution. Hence, we assumed constant genetic parameters for the logarithm of *R*_0_. The same approach has been used to model genetic variation in the residual variance, which is also restricted to non-negative values (SanCristobal-Gaudy *et al*. 1998; Hill and Mulder 2010). The log-normal distribution of genotypic values for *R*_0_ results in a decrease of the genetic standard deviation in *R*_0_ with decreasing *R*_0_ (Figure 2), which seems reasonable given the presence of a lower bound for *R*_0_. Moreover, the log-normal distribution for *R*_0_ is convenient, because it results in simple expressions for the breeding value and the genetic variance for prevalence.

The assumption of a normal distribution for the logarithm of genotypic values for *R*_0_ also agrees with the standard implementation of generalized linear (mixed) models (GLMM; Nelder and Wedderburn, 1972). *R*_0_ refers to an expected number of infected individuals; In other words, *R*_0_ is the expected value of count data. In GLMM, the default link function for count data is the log-link (McCullagh and Nelder, 2019). Hence, our linear model for the logarithm of *R*_0_ also agrees with common statistical practise.

The strong increase of the genetic variance in prevalence with decreasing *R*_0_ (Figure 5A) is not due to the assumption of lognormality of *R*_0_. On the contrary, the log-normal distribution results in a decrease of the genetic standard deviation in *R*_0_ with decreasing *R*_0_ (Figure 2). The strong increase in the genetic variance in prevalence results from the relationship between *R*_0_ and the prevalence in the endemic steady state (Figure 1; Equation 3), which becomes steeper when *R*_0_ is closer to one. This relationship is very well established in epidemiology since Weiss and Dishon (1971; *e*.*g*., Keeling & Rohani, 2011).

While we defined an additive genetic model for the logarithm of *R*_0_, we can also find the additive genetic effect (breeding value) for *R*_0_ itself. Using results of Appendix 4,

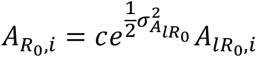

Hence, the 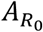 represents the additive component of the genotypic value for *R*_0_. However, because our model is additive on the log-scale, while the genotypic value for *R*_0_ includes non-additive genetic effects, we decided to build our theory on the breeding value for the logarithm of *R*_0_.

### Other compartmental models

In this work, we focused on endemic infectious diseases following a SIS-model, where individuals can be either susceptible (S, *i*.*e*., non-infected) or infected (I). Hence, we assumed the infection does not confer any long-lasting immunity, and we ignored the potential existence of infected classes (“compartments”) other than S and I, such as recovered infected individuals that are not yet susceptible again. Moreover, we ignored the influx of new individuals into the population due to births, and the removal of individuals due to deaths.

A key condition for validity of our results is that the pathogen can replicate only in the host individual, meaning that a reduction in infected individuals fully translates into reduced exposure of the host population to the pathogen. The mere survival of the pathogen in the environment does not violate our assumptions; see Hulst *et al*., 2021 for a discussion. Our conclusions are not limited to SIS models if this condition is met, but apply to all models with no longer lasting immunity. For models with temporary immunity (*e*.*g*., SIRS) or lifelong immunity (*e*.*g*., SIR) the conclusions with respect to infectivity and susceptibility will be true, but the genetic variation in recovery may play no role (*e*.*g*., SIR) or a different more restricted role (*e*.*g*., SIRS when the R compartment lasts long).

Also infections that do confer long-lasting immunity may show endemic behavior when a population is large enough. Measles in the human population before the introduction of vaccination are is a well-known example. For such infections, the same mechanisms as discussed above will play a role and the endemic prevalence for a homogeneous population still follows from Equation 3. However, the introduction of new susceptibles by birth can no longer be ignored, and recovery of infected individuals does not result in new susceptible individuals. Thus the role of recovery will change, and the genetic make-up of the newborn individuals becomes relevant, particularly in populations undergoing selection.

### Positive feedback

The increasing difference between the breeding value for prevalence and the breeding value for individual infection status at lower prevalence (Equation 33) is a result of the increasing slope of the relationship between *R*_0_ and the endemic prevalence (Equation 3, Figure 1). Equation 3 follows directly from a simple equilibrium condition (see text above Equation 3). However, the focus on the equilibrium partly obscures the underlying mechanism. Figures 10A and B illustrate that the difference between *A*_*P*_ and *A*_*y*_ originates from positive feedback effects in the transmission dynamics. (Figure 10 shows results for selection against susceptibility, selection for faster recovery would yield identical results). With lower susceptibility fewer individuals will become infected, which subsequently translates into a reduced transmission rate, followed by a further reduction in the number of infected individuals, *etc*, resulting in a positive feedback loop (Figure 10A). The initial change in prevalence before feedback effects manifest is equal to the selection differential in breeding value for individual infection status (ΔĀ_*y*_; horizontal lines in Figure 9). This change represents the direct response due to reduced susceptibility, and does not include any change in exposure of susceptible individuals to infected herd mates. Next, prevalence decreases further because the initial decrease in prevalence reduces the exposure of susceptible individuals to infected herd mates. This additional decrease represents the indirect response to selection via the “social” environment. Without genetic variation in infectivity, the direct response makes up a fraction *P* of the total response in prevalence, and the indirect response a fraction 1 − *P*.

**Figure 10.**
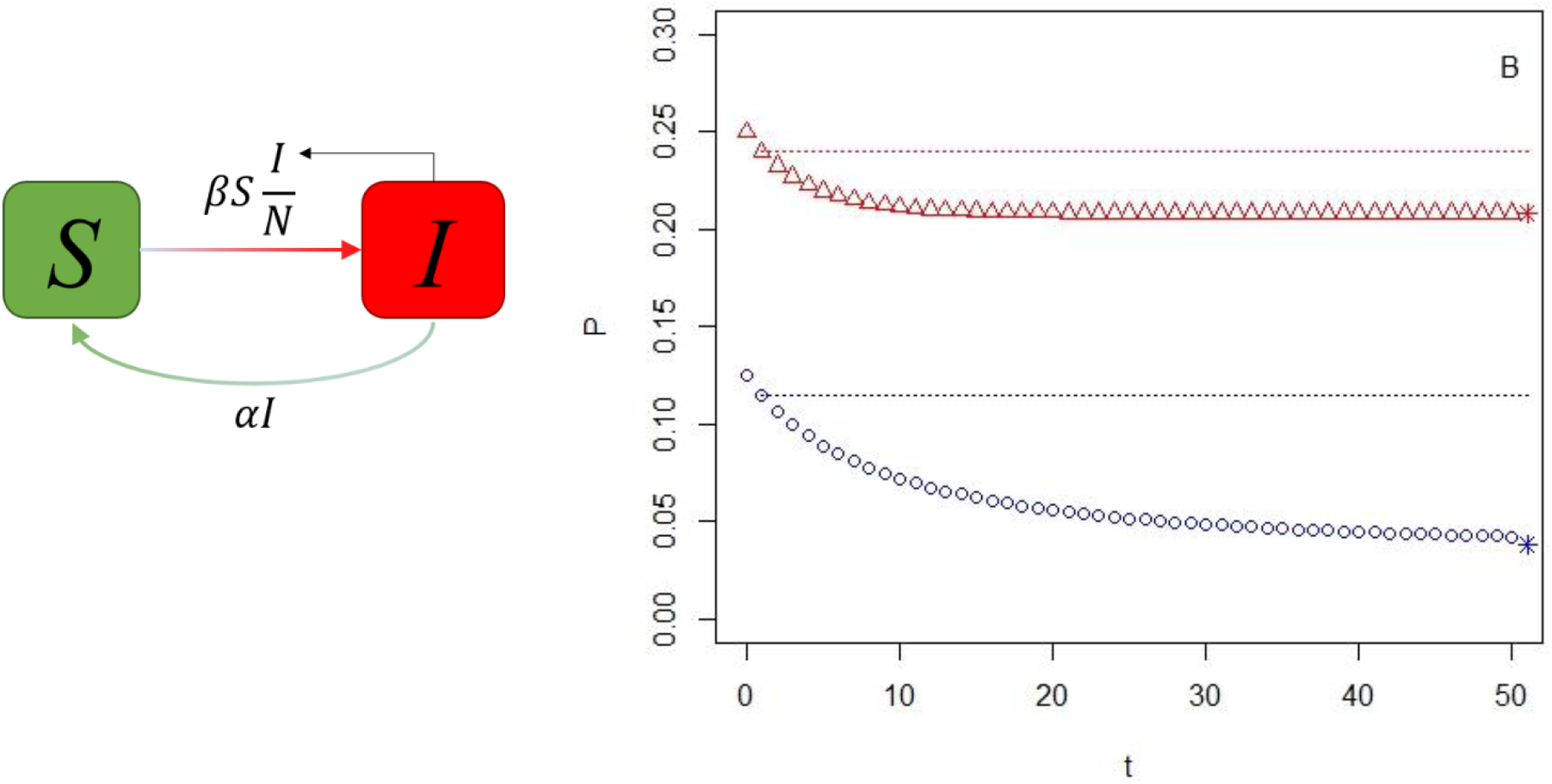
Positive feedback after selection for lower susceptibility. Panel A: Diagram of the SIS compartmental model illustrating the feedback, with the number of susceptible (*S*) and infectious (*I*) individuals and the transmission and recovery rates (ignoring heterogeneity for simplicity). A reduction in the transmission rate parameter *β* reduces *I*, which in turn reduces the transmission rate, leading to a further reduction in *I, etc*. Panel B: Convergence of the prevalence to the new equilibrium after selection. For two populations; one starting at *P* = 0.25 (red triangles; *c* = 1.333), the other at *P* = 0.125 (blue circles; *c* = 1.143). The *x*-axis represents cycles of the transmission loop. The horizontal dotted lines show the prevalence predicted by the breeding value for binary infection status, and represent the direct effect. The Asterix shows the equilibrium prevalence after convergence, which occurs a little later than *t* = 50 for the lower line. The genetic selection differential for binary infection status equals ΔĀ_*y*_ = −0.01 for each of the two populations. The initial response to selection (the *y*-axis difference between *t* = 0 and *t* = 1) is equal to the ΔĀ_*y*_ of −0.01 for both scenarios. Total response is −0.04 for the scenario with *P* = 0.25, and −0.08 for the scenario with *P* = 0.125, corresponding to −0.01/0.25 and −0.01/0.125. Results in panel B follow from iterating on Equation 20a, using a single value for ℛ_0,*i*_, with 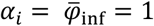 and choosing γ so that the selection differential Δ*Ā*_*y*_ = −0.01 (using 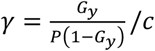 from Equation 20a). In each iteration, the *P* in the righthand side of Equation 20a is replaced by the *P*_*i*_ calculated from Equation 20a in the previous iteration. This iteration converges to the prevalence given by Equation 3 (assuming negligible heterogeneity).

### Herd immunity

In Figure 8A, the infection ultimately goes extinct due to mass selection for individual infection status. This happens due to a phenomenon known as herd immunity (Fine, 1993). In the final generation, the infection disappears because *R*_0_ falls below a value of one; not because all the individuals have become fully resistant to infection. This result is similar to the eradication of an infection by means of vaccination, which also does not require full immunity of all individuals and can also be achieved when only part of a population is vaccinated (Anderson and May, 1985). As can be seen in Figure 10 and in simulation results of Hulst *et al*. (2021), herd immunity develops over cycles of the transmission-recovery loop. Thus the full benefits of genetic selection or vaccination do not manifest immediately, as it takes some time for a population to converge to the new endemic steady state.

The relevance of herd immunity for response to genetic selection can be illustrated using the data underlying Figure 8A. For the population starting at a prevalence of 0.5, the contact rate is equal to two, and the mean breeding value for log-susceptibility is equal to zero in the initial generation (*c* = 2, Ā_*l*γ_ = 0, so that 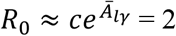). In the final generation, the mean breeding value for log-susceptibility has dropped to −0.73, so that *R*_0_ ≈ 2*e*^−0.73^ = 0.96. Hence, *R*_0_ < 1, explaining extinction. However, if the average individual of the final generation would have been exposed to the infection pressure of the first generation, then the expected prevalence for this individual would have been 0.32 (From Equation 20a, with *R*_0.*i*_ = 0.96 and *P* = 0.5). Hence, the individual would have been infected 32% of the time. Nevertheless, in a population consisting entirely of this type of individual, as is the case in the final generation, the infection will no longer be present in the long term. This example illustrates the relevance of indirect effects for herd immunity and for response to selection of infectious diseases.

### Utilization of hidden genetic variation for genetic improvement

In this work, we have shown that a fraction 1 − *P* of the full individual genetic effect on the endemic prevalence represents an IGE, because only a fraction *P* of the full effect surfaces in the infection status of the individual itself (excluding genetic variation in infectivity; Equation 33 and Appendix 5). In other words, a fraction 1 − *P* of the individual genetic effects of susceptibility and recovery on the prevalence are hidden to direct selection and classical genetic analysis. Nevertheless, results in Figure 8 show that prevalence responds rapidly to selection, particularly when prevalence is small. Hence, prevalence responds faster to selection when a greater proportion of its heritable variation is hidden, and when heritability is low (Figure 7A), which seems a paradox.

However, the IGEs due to susceptibility and recovery are a special kind, because they are fully correlated to the corresponding DGE. For each of the two traits, there is only a single genetic effect (*A*_*l*γ_ and *A*_*lα*_, respectively), which has both a direct effect and an indirect effect on the prevalence. Hence, when selection changes the mean DGE, the mean IGE changes correspondingly. This can be seen from Equation 38, where the term 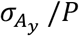 represents the full additive genetic standard deviation in prevalence (as is clear from Equation 33), while the accuracy 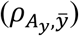 refers to selection for the direct effect only. Hence, without genetic variation in infectivity, the total response to selection based on individual infection status can be interpreted as the sum of a direct response in DGE and a correlated response in IGE,

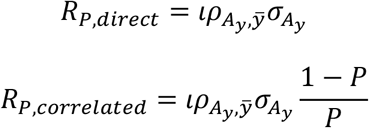

and the sum of *R*_*P,direct*_ and *R*_*P,correlated*_ is equal to Equation 38. The *R*_*P,direct*_ is the expected response to selection based on ordinary genetic analysis of individual infection status. The *R*_*P,correlated*_ represents the additional response due to IGE. The direct response occurs immediately in the first cycle of the transmission loop (Figure 10B), while the indirect response manifests gradually over several cycles of the transmission-recovery loop, particularly when prevalence is small (see also result in Hulst *et al*. 2021).

The response due to the IGE of susceptibility and recovery arises naturally when selecting for lower individual infection status (*i*.*e*., for the direct effect); it does not require any specific measures of the breeder. Thus, on the one hand, our results imply that response to genetic selection against infectious diseases should be considerably greater than currently believed, even when no changes are made to the selection strategy. While empirical studies are scarce, the available results support this expectation (discussed in Hulst *et al*. 2021).

On the other hand, however, classical selection for direct effects is not the optimal way to reduce prevalence, for the following two reasons. First, classical selection does not target genetic effects on infectivity, because an individual’s infectivity does not affect its own infection status (Lipschutz-Powell et al., 2012). Hence, infectivity changes merely due to a potential genetic correlation with susceptibility and/or recovery. When this correlation is unfavourable, infectivity will increase and response in prevalence will be smaller than expected based on the genetic selection differentials for susceptibility and recovery. (And thus smaller than the result of Equation 38). In theory, this could even lead to a negative net response (Griffing 1967). This is similar to the case with social behaviour-related IGEs on survival in laying hens and Japanese quail, where selection for individual survival has sometimes increased mortality (Craig and Muir, 1996; Muir 2005). This scenario seems unlikely for infectious diseases, but at present we lack knowledge of the multivariate genetic parameters of susceptibility, infectivity and recovery to make well-founded statements.

Second, even in the absence of genetic variation in infectivity, individual selection for susceptibility and recovery is non-optimal because the accuracy of selection is limited due to limited heritability, particularly at low prevalence (Figure 7A). The response to selection in traits affected by IGE can be increased by using kin selection and/or group selection (Griffing 1976; Muir 1996; Bijma 2011), or by including IGE in the genetic analysis (Muir 2005, Bijma *et al*. 2007b; Muir *et al*. 2013; Biemans *et al*. 2019, Anacleto *et al*. 2015, Pooley *et al*. 2020). Kin selection occurs when transmission takes place between related individuals, for example within groups of relatives (Anche *et al*. 2014). Group selection refers to the selection of parents for the next generation based on the prevalence in the group in which transmission takes place, rather than on individual infection status (Griffing 1976). Both theoretical and empirical work shows that kin and group selection lead to utilization of the full genetic variation, including both DGE and IGE (Griffing 1976, Muir 1996, 2005; Bijma and Wade, 2008; Bijma 2010, 2011). For infectious diseases, the work of Anche *et al*. (2014) illustrates the effect of kin selection, where favourable alleles for susceptibility increase much faster in frequency when disease transmission is between related individuals. Simulation studies on IGE in pig populations suggest that the benefits of kin selection should also extend to breeding schemes based on genomic prediction (Chu *et al*. 2021).

### Do pathogens create kin selection?

Exposure to infectious pathogens is a major driver of the evolution of host populations by natural selection, both in animals and plants (reviewed in Karlsson *et al*. 2014 and Ebert and Fields 2020). In the human species, for example, a study of genetic variation in 50 worldwide populations reveals that selection on infectious pathogens is the primary driver of local adaptation and the strongest selective force that shapes the human genome (Barreiro and Quintana-Murci 2010; Fumagalli *et al*. 2011). The key role of infectious pathogens in natural selection, together with the large contribution of IGE to the genetic variation in prevalence in the host population, indicates that IGE must have been an important fitness component in the evolutionary history of populations. This, in turn, suggests that associating with kin may have evolved as an adaptive behaviour. In other words, the key role of infectious diseases in natural selection might lead to social structures where individuals associate preferably with kin, because such behaviour has indirect fitness benefits. This is because interactions among kin lead to utilisation of the full heritable variation in fitness, including both DGE and IGE (Bijma, 2010), and thus accelerate response of fitness to selection. At low to moderate levels of the endemic prevalence the indirect genetic variance in prevalence might be sufficiently large for such behaviour to evolve, even in the absence of direct fitness benefits such as preferential behaviour towards kin. While this is a complex issue requiring careful quantitative modelling, including migration and the emergence of selfish mutants, the key role of pathogens in natural selection together with the large IGE demonstrated here strongly suggest the importance of kin selection in the history of life.

In agriculture, the implementation of kin selection may be feasible when animals can be kept in kin groups or plants can be grown in plots of a single genotype or a family in the breeding population. In many cases, however, this will not be feasible, and other methods are required to optimally capture the IGE underlying the prevalence of infectious diseases. Current developments in sensing technology and artificial intelligence enable the development of tools for large scale automated collection of longitudinal data on individual infection status, and also on the contact structure between individuals (relevant mainly in animals). These advances, together with genomic prediction and recently developed statistical methods for the estimation of the direct and indirect genetic effects underlying the transmission of the infection (Biemans *et al*., 2019; Pooley *et al*. 2020), could represent a much-needed breakthrough in artificial selection against infectious diseases in agriculture. Our results on genetic variation and response to selection suggest that such selection is way more promising than currently believed.

## Appendix 1

### *R*_0_ with heterogeneity and log-normally distributed susceptibility, infectivity en recovery

We assume that the transmission rate from infected individual *j* to susceptible individual *i* is proportional to the product of the infectivity of *j* and the susceptibility of *i* (Equation 5),

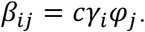

So there is no interaction between *i* and *j*. (This property is known as separable mixing in the epidemiological literature; Diekmann *et al*. 1990; 2013). Moreover, we assume that susceptibility, infectivity and recovery follow a log-normal distribution (Equations 6 and 7). We also assume that the population is not very small, so that in the early phase of an endemic where only few individuals are infected, the composition of the remaining susceptible individuals is not affected.

Because *R*_0_ refers to the “total number of individuals that become infected by a typical infected individual over its entire infectious lifetime”, we define an individual lifetime infectivity, which is the product of an individual’s infectivity per unit of time and the average duration of its infectious lifetime,

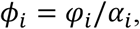

which follows a log-normal distribution with parameters following from those of *φ* and *α*. Hence, we have condensed our three genetic effects into two.

We can find *R*_0_ from

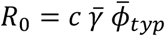

where 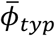 is the lifetime infectivity of the typical infected individual, and 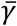 is the simple average susceptibility in the population,

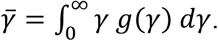

where *g*(γ) is the pdf of γ. We can use the simple average of susceptibility in this expression because we assume the population is large.

With separable mixing, the typical infected individual is created immediately in the first generation of disease transmission. This is the case because there is no interaction between γ and *φ*, so that the properties of the typical infected individual are determined entirely by susceptibility. Hence, the pdf of γ for the typical infected individual follows from weighing *g*(γ) by γ,

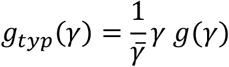

Since the properties of the typical infected individual depend on susceptibility only, we can find 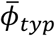 by averaging *ϕ* over its distribution conditional on γ, and subsequently averaging over the distribution of γ,

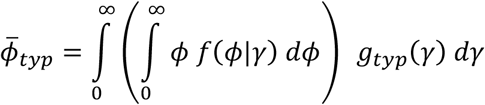

Hence, we now have the elements of *R*_0_, but still need to solve the integral expression.

Because conditional Normal distributions are also Normal and the logarithm is a bijective function, *ϕ*|γ follows a log-normal distribution with parameters being the conditional mean and variance of the Normal distribution,

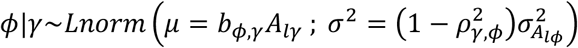

with *b*_*ϕ*,γ_ = *cov*(*A*_*l*γ_, *A*_*lϕ*_)/*var*(*A*_*l*γ_) denoting the regression coefficient of *A*_*lϕ*_ on *A*_*l*γ_, and 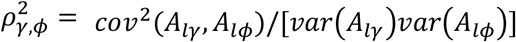 the squared correlation, where *A*_*lϕ*_ denotes the breeding value for logarithm of lifetime infectivity.

Hence, the inner integral is the mean of a log-normal variate, which is of the form 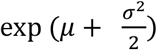

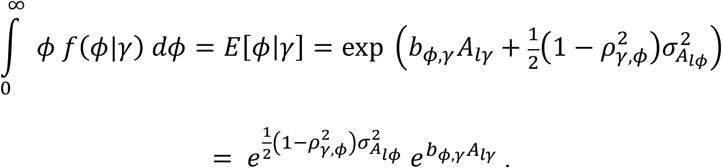

Since the first term of this expression is a constant,

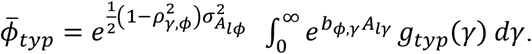

Substituting 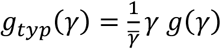, and replacing *g*(γ) by the corresponding log-normal pdf yields

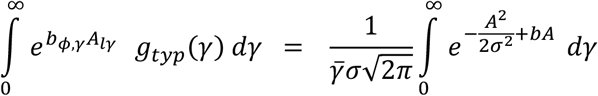

where we simplified the notation for brevity, using 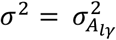, and *A* = *A*_*l*γ_. Next, we change variable, using *d*γ = *e* ^*A*^*dA*, and adjust the bounds accordingly,

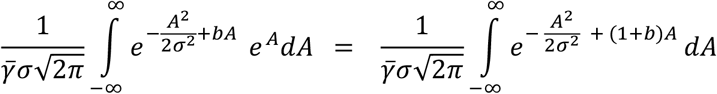

Solving the integral term in Mathematica-online yields

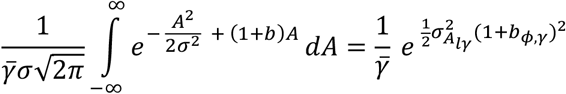

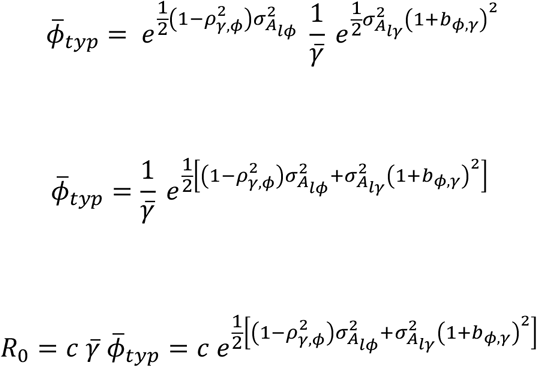

Using 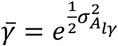 and 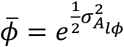 this simplifies to Equations 14 and 16 of the main text,

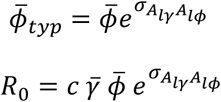

Further simplification follows from expressing 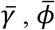 and 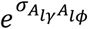 in terms of variances and covariances of γ, *φ* and *α*.

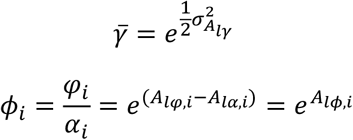

where *A*_*lϕ,i*_ = *A*_*lφ,i*_ − *A*_*lα,i*_, which is the breeding value for the logarithm of lifetime infectivity, with

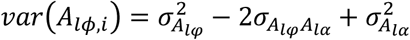

From the properties of the log-normal distribution,

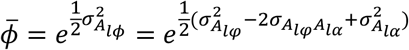

Furthermore,

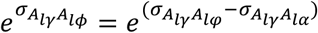

Substitution of the expressions for 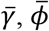 and *e*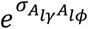 into 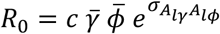 yields

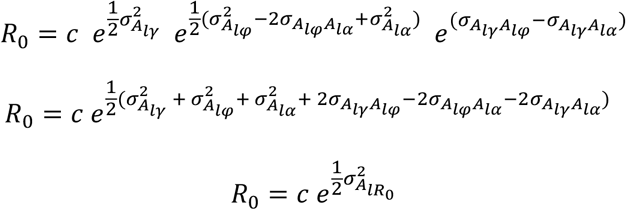

The right-hand side of this expression is identical to the mean genotypic value for *R*_0_ (Equation 12).

## Appendix 2

### Numerical solution to find the endemic equilibrium prevalence with heterogeneity

To find the endemic prevalence, *P*, we partition the population into types, *i*, and numerically solve the expressions

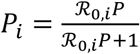

and

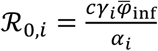

for *P*. Here we develop this numerical solution for the case where susceptibility, infectivity and recovery follow a log-normal distribution, assuming separable mixing (see Appendix 1).

As can be seen from the expression for ℛ_0,*i*_, the equilibrium prevalence for a type depends on both its susceptibility (γ_*i*_) and its recovery rate (*α*_*i*_). Individuals with above-average susceptibility are over-represented among the infecteds in the equilibrium, whereas individuals with above-average recovery are under-represented. Hence, as can be seen from the expression for ℛ_0,*i*_, the partitioning into types should be based on γ_*i*_/*α*_*i*_. Therefore we define

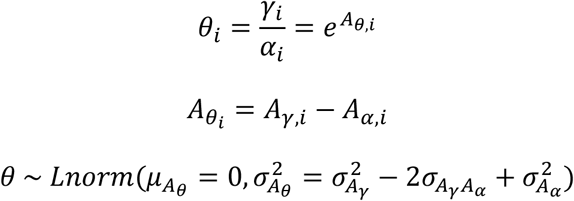

To numerically solve the two equations given above, we also need 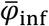. The 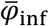 will depends on the *θ*_*i*_ of the infecteds when infectivity is correlated to susceptibility and/or recovery. Hence, we need the distribution of *φ*|*θ*, which follows from

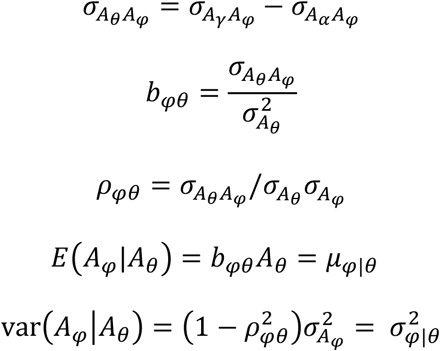

so that

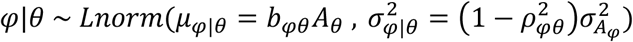

From the log-normal distribution:

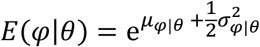

Hence, we can partitioning *θ* into classes *i*, with

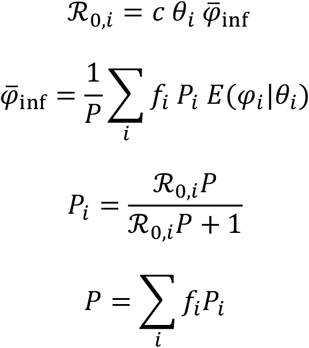

where *f*_*i*_ is the fraction of individuals of type *i, f*_*i*_ = *N*_*i*_/*N*, and *P*_*i*_ is the prevalence in type *i, P*_*i*_ = *I*_*i*_/*N*_*i*_. The numerical solution follows from iterating on these four equations. An R-code is in Supplementary Material 1.

## Appendix 3

### Methods for simulation of epidemics and validation of prevalence and genotypic value for individual disease status

We simulated endemics according to standard epidemiological theory to validate the numerical solution of the endemic prevalence (Equations 20a,b) and the genotypic values for binary disease status (Equation 24). We considered two compartments of individuals, susceptible individuals (S) and infected (I) individuals, and a so-called stochastic SIS-model where susceptible individuals can become infected, and infected individuals can recover and then immediately become susceptible again (Weiss and Dishon, 1971). For simplicity, we simulated genetic variation in susceptibility only, with 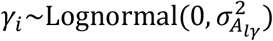.

To limit Monte-Carlo error, we simulated a relatively large population of *N* = 2,000 genetically unrelated individuals for a total of 300,000 events (infection or recovery). We used a burn-in of 100,000 events before recording data on individual binary disease status. Hence, in the recorded data, the average individual experienced 100 events (50 infections and 50 recoveries).

The endemic was started by infecting a proportion *P*_0_ = 1-1/*c* of the individuals, chosen at random. Subsequently, we sampled events (infection or recovery) and the individual involved using Gillespie’s algorithm (Gillespie, 1977). For each infected individual, the probability of recovery was proportional to the recovery rate, *α*. For susceptible individual *i* the probability of infection was proportional to *c*γ_*i*_*I*/*N, I*/*N* denoting the fraction of the population that is infected. Probabilities were accumulated over all individuals and scaled to a sum of 1 by dividing them by their sum. Finally, the specific event was sampled by drawing a random number, say *x*, from a standard uniform distribution and finding the event and the corresponding individual belonging to the probability interval [*x*_*l*_, *x*_*h*_], where *x*_*l*_ < *x* < *x*_*h*_. The disease status of that individual and *I* were updated before sampling the next event. The time of each event was not simulated. After 300,000 events, prevalence was calculated as the disease status averaged over the entire population, and also by individual, discarding the burn-in period. The regression coefficient of average individual disease status on *G*_*y*_ was also estimated.

Additive genetic variance in log-susceptibility was 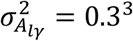. Three scenarios were considered, differing in contact rate: *c* = 1.22 giving *P* = 0.2, *c* = 2 giving *P* = 5 and *c* = 5.15 giving *P* = 0.8. Those combinations of 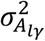, *c* and *P* were found by numerically solving Equations 20a&b. The actual prevalences observed in the simulations were equal to these numerical solutions.

## Appendix 4.

### Additive genetic variance in log-normal traits

We assumed log-normally distributed genotypic values for susceptibility, infectivity and recovery, also resulting in a log-normal distribution for 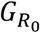 and for 1 − *G*_*P*_. Hence, genetic effects are additive on the log-scale, but taking the exponent introduces some non-additive genetic variance on the actual scale. Here we derive the fraction of the variance that is additive on the actual scale.

Because all genetic effects had a mean of zero on the log-scale, the problem is equivalent to finding the fraction of additive variance in *z* = *e*^*x*^, where *x*∼*N*(*μ* = 0, *σ*^2^). From the properties of the log-normal distribution, 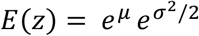. With a small change *dμ*, the mean of *z* becomes 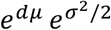. Hence, the mean of *z* changes by an amount 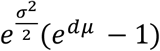. Since 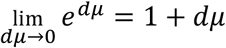, this change corresponds to 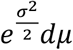. Thus the least-squares linear regression coefficient of *z* on *x* equals

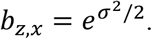

For example, the linear regression coefficient of the genotypic value for prevalence (Equation 26b) on the breeding value for the logarithm of *R*_0_ equals 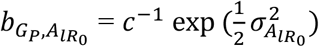. Thus the additive effect for *z* equals

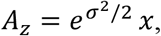

and additive variance in *z* equals

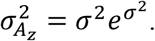

The total variance in *z* follows from the properties of the log-normal distribution,

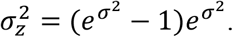

The additive fraction of the variance in *z*, therefore, equals

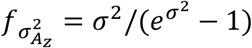

Figure A3.1 illustrates that the additive fraction of the variance in *z* approaches 1 when *σ*^2^ goes to zero. For *σ*^2^ = 0.5^2^, ∼88% of the variance in *z* is additive. Variances on the log scale larger than 0.5^2^ are unrealistic (see main text). This indicates that at least 88% of the genetic variance in susceptibility, infectivity, recovery, *R*_0_ and prevalence is additive when they follow a log-normal distribution.

**Figure A4.1.**
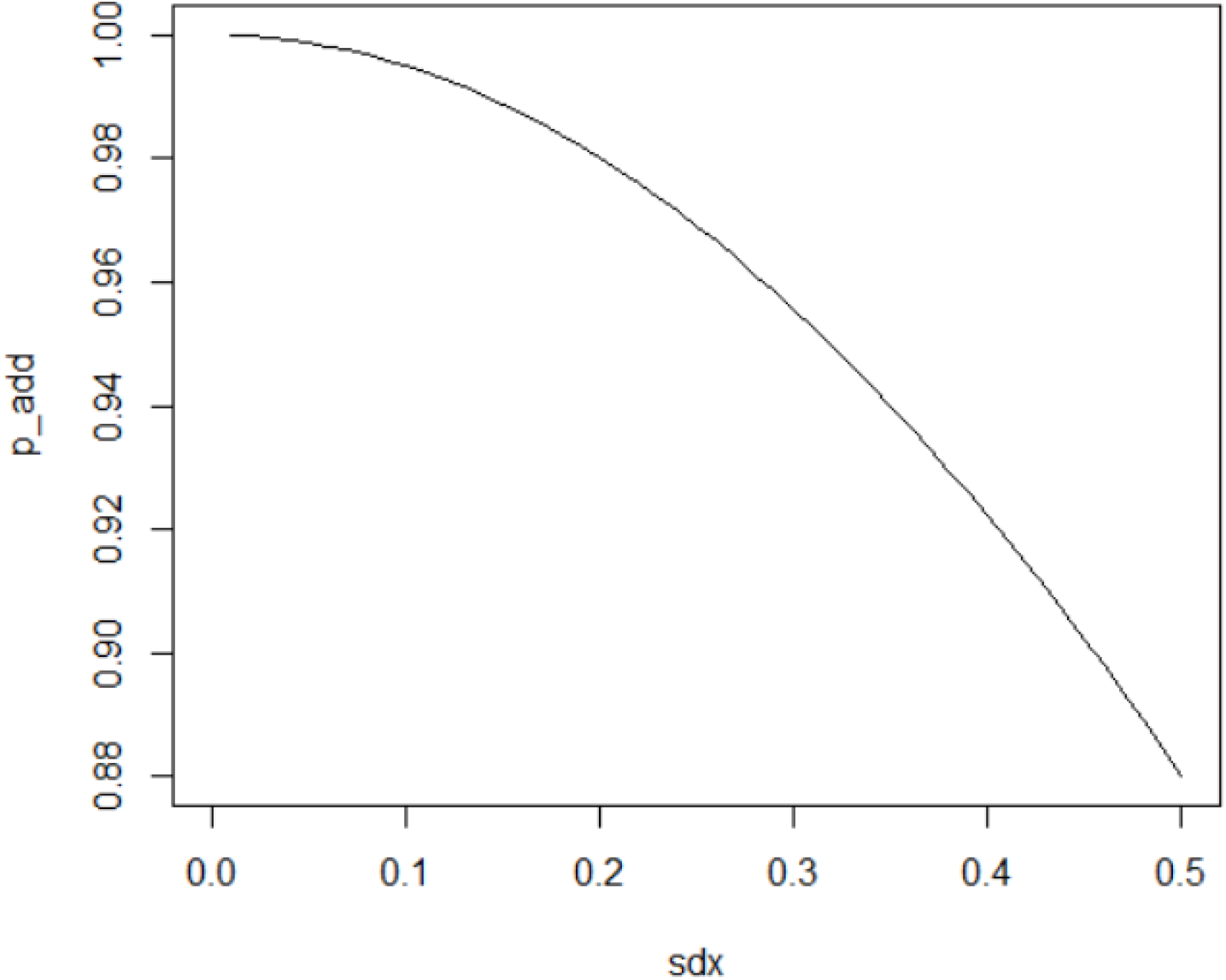
The additive fraction of the variance in traits following a log-normal distribution. sdx denotes the standard deviation on the normal scale.

## Appendix 5.

### Breeding value for individual disease status *vs*. breeding value for prevalence, without genetic variation in infectivity

Without genetic variation in infectivity we have *φ*_*i*_ = *φ* = 1, because the scale is included in the effective contact rate *c*. From Equation 24, the genotypic value for individual binary disease status is,

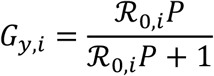

where, from Equation 20b,

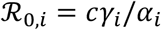

From Equation 26a, the genotypic value for prevalence is,

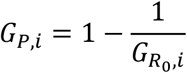

where, from Equation 8,

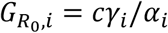

Hence, without genetic variation in infectivity, ℛ_0,*i*_ and 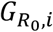 are identical, and we will use the symbol 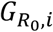 in the following.

The linear approximation of the relationship between *G*_*y*_ and *G*_*P*_ follows from a comparison of their first derivatives with respect to 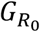,

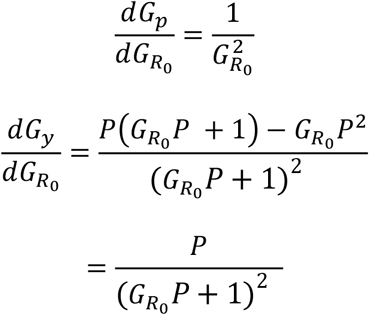

Substituting Equation 3, assuming limited heterogeneity, yields

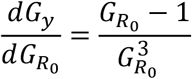

Hence,

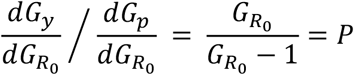

Therefore, for a small change in an individual’s genotypic value for *R*_0_, the change in its genotypic value for binary disease status is only a fraction *P* of the change in its genotypic value for prevalence,

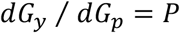

Hence, when expressed relative to their mean, *G*_*y*_ and *G*_*p*_ differ approximately by a factor *P* (see also Figure 4 in Bijma 2020). This result is approximate, because the true relationship is non-linear and the expression *P*_*eq*_ = 1 − 1/*R*_0_ is approximate with variation among individuals. For realistic magnitudes of the genetic variance, however, the non-linearity is limited. Note that the above derivation does not require the assumption of a log-normal distribution of susceptibility and recovery.

So far, this appendix has considered genotypic values. However, the factor *P* also applies on the level of the breeding values, which can be shown as follows. The above derivation is based on derivatives with respect to genotypic value for *R*_0_, say 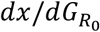, where *x* is *G*_*y*_ or *G*_*P*_. Since the relationship between *P* an *y* arises entirely via 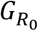, we can translate the results to the breeding values using

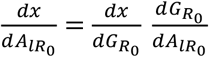

The latter, 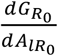, is the same for *x* is *G*_y_ or *x* is *G*_*P*_, so that the ratio *dA*_*p*_ ⁄*dA*_*p*_ is the same as *dG*_*y*_⁄ *dG*_*p*_. Hence, we also find

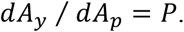

## Appendix 6

### Methods for observed response to selection

First a base population was generated of *N* = 4000 unrelated individuals, with genetic variation in susceptibility only. No distinction was made between males and females. For each individual, breeding values for the logarithm of susceptibility were sampled from *A*_*l*γ_∼*N*(0, 0.3^2^), and individual susceptibility was calculated as 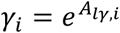. The expected prevalence for the base generation was calculated as *P*_0_ = 1 − 1/*c*, with a *c* of either 2 or 10, and the initial disease status of base generation individuals was sampled at random from *Bernoulli*(*P*_0_).

Next, an endemic was simulated following methods described in Appendix 3, for a total of 15,000 events (sum of infections and recoveries), consisting of a burn-in of 10,000 events and 5,000 recorded events. The 4,000 individuals were ordered based on their mean individual disease status over the 5,000 recorded events (so based on 1.25 events on average per individual), and the 2000 individuals with the lowest values were selected as parents of the next generation (corresponding to a selected proportion of 0.5).

Selected parents were mated at random. Each pair of parents produced two offspring, resulting in *N* = 4,000 offspring. Offspring inherited the breeding value for the logarithm of susceptibility in a Mendelian fashion; 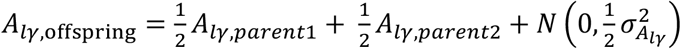. The initial disease status of offspring (*i*.*e*., at the start of the burn-in period of their generation) was sampled at random from *Bernoulli*(*P*_offspring_), where *P*_offspring_ denotes the expected prevalence in the offspring generation, calculated as 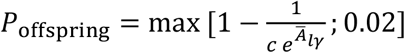. The 0.02 guaranteed an average of at least 80 infected individuals at the start of the endemic in any generation, also when the expected prevalence was zero (*i.e*, when 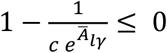. Then an endemic was started, as described above for the base generation, *etc*. This process was repeated until the number of infected individuals dropped to zero, implying extinction of the infection.

## Supplementary material

**Supplementary Material 1:** R-code to numerically find the endemic equilibrium prevalence. See file “Supplementary Material 1 - Numerical Solution Prevalence Heterogeneity.R”

**Supplementary Material 2:** Additional results for the endemic equilibrium prevalence. See file “Supplementary Material 2 - Effect of heterogeneity on endemic prevalence.xlsx”. Note that, when *c* < 2, prevalence is (much) higher than 1 − 1/*c*, while prevalence is only a little lower than 1 − 1/*c* when *c* > 2. Hence, for *c* > 2, *P* = 1 − 1/*c* is a useful predictor.

**Supplementary Material 3:** Validation of Equation 34 Equation 34 states that, in the absence of genetic variation in infectivity, the breeding value for individual disease status is given by

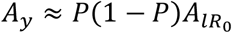

where *P* denotes the endemic prevalence and 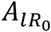 the breeding value for the logarithm om *R*_0_. We validated this expression using stochastic simulation of endemics following standard epidemiological theory (as outlined in Appendix 3). We simulated a population of *N* = 10,000 individuals, for a total of 500,000 events (sum of infections and recoveries), using the first 100,000 events as burn in. We considered input values of 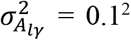 0.3^2^ and 0.5^2^. We did not simulate genetic variation in the recovery rate, because this has the identical effect as genetic variation in susceptibility (Equations 21 and 22). We also did not simulate genetic variation in infectivity, since the expression we intend to validate here refers to the case without genetic variation in infectivity (In other words, the infectivity component has to be left out of the 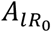 in Equation 34). We considered a prevalence ranging from 0.1 to 0.9, with steps of 0.1, and found the values for the contact rate that correspond to these prevalences numerically from solving Equations 20a and b (Table S3.1; R-code in Supplementary Material 1).

**Table 3.1.**
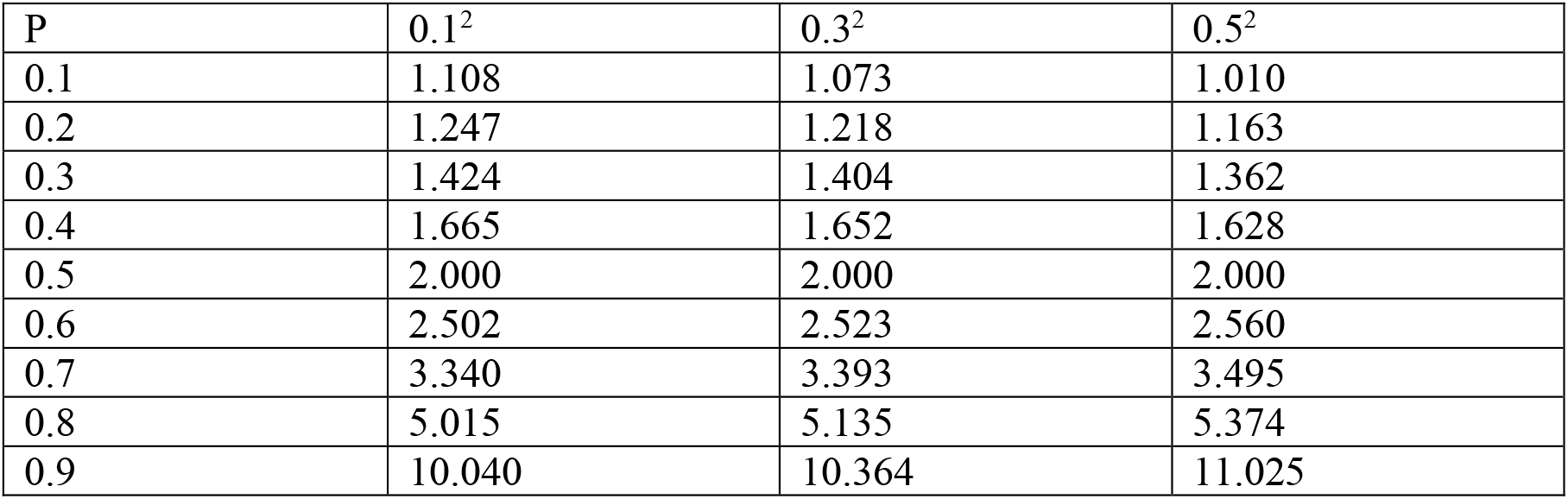
Contact rates required to find a certain prevalence, for 3 values of 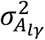.

By definition, the individual breeding value equals the regression of individual trait value on additive genetic effects. Validation, therefore, focussed on the comparison of the *P*(1 − *P*) term in the above expression for *A*_*y*_ to the regression coefficient of individual disease status on 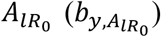 estimated from the simulated data. Table S3.2 shows close agreement between these two parameters.

**Table S3.2.** Comparison of estimated regression coefficient (bhat) of individual disease status on individual breeding value for the logarithm of *R*_0_ with *P*(1-*P*). For three levels of genetic variance in log(R_0_). P_desired denotes the desired prevalence; P_realized denotes the realized prevalence obtained using the contact rates given in Table S3.1.

|  | v(A_log_R0) |  |  | v(A_log_R0) |  |  | v(A_log_R0) |  |  |
| --- | --- | --- | --- | --- | --- | --- | --- | --- | --- |
|  | 0.1^2 | 0.1^2 | 0.1^2 | 0.3^2 | 0.3^2 | 0.3^2 | 0.5^2 | 0.5^2 | 0.5^2 |
| P_desired | P_realized | bhat | P(1-P) | P_realized | bhat | P(1-P) | P_realized | bhat | P(1-P) |
| 0.10 | 0.10 | 0.09 | 0.09 | 0.10 | 0.09 | 0.09 | 0.11 | 0.10 | 0.10 |
| 0.20 | 0.20 | 0.16 | 0.16 | 0.20 | 0.16 | 0.16 | 0.20 | 0.16 | 0.16 |
| 0.30 | 0.30 | 0.21 | 0.21 | 0.31 | 0.21 | 0.21 | 0.30 | 0.20 | 0.21 |
| 0.40 | 0.40 | 0.25 | 0.24 | 0.40 | 0.23 | 0.24 | 0.40 | 0.23 | 0.24 |
| 0.50 | 0.50 | 0.25 | 0.25 | 0.50 | 0.24 | 0.25 | 0.50 | 0.24 | 0.25 |
| 0.60 | 0.60 | 0.23 | 0.24 | 0.60 | 0.23 | 0.24 | 0.60 | 0.23 | 0.24 |
| 0.70 | 0.70 | 0.21 | 0.21 | 0.70 | 0.21 | 0.21 | 0.70 | 0.20 | 0.21 |
| 0.80 | 0.80 | 0.16 | 0.16 | 0.80 | 0.16 | 0.16 | 0.80 | 0.15 | 0.16 |
| 0.90 | 0.90 | 0.09 | 0.09 | 0.90 | 0.09 | 0.09 | 0.90 | 0.09 | 0.09 |

